# A tale of two n-backs: Diverging associations of dorsolateral prefrontal cortex activation with n-back task performance

**DOI:** 10.1101/2024.05.23.595597

**Authors:** Philip N. Tubiolo, John C. Williams, Jared X. Van Snellenberg

**Author notes:** CORRESPONDING AUTHOR: Jared X. Van Snellenberg, 101 Nicolls Rd, Health Sciences Center 10-087J, Stony Brook, NY 11794.

## Abstract

**Background:** In studying the neural correlates of working memory (WM) ability via functional magnetic resonance imaging (fMRI) in health and disease, it is relatively uncommon for investigators to report associations between brain activation and measures of task performance. Additionally, how the choice of WM task impacts observed activation-performance relationships is poorly understood. We sought to illustrate the impact of WM task on brain-behavior correlations using two large, publicly available datasets.

**Methods:** We conducted between-participants analyses of task-based fMRI data from two publicly available datasets: the Human Connectome Project (HCP; n = 866) and the Queensland Twin Imaging (QTIM) Study (n = 459). Participants performed two distinct variations of the *n-*back WM task with different stimuli, timings, and response paradigms. Associations between brain activation ([2-back − 0-back] contrast) and task performance (2-back % correct) were investigated separately in each dataset, as well as across datasets, within the dorsolateral prefrontal cortex (dlPFC), medial prefrontal cortex, and whole cortex.

**Results:** Global patterns of activation to task were similar in both datasets. However, opposite associations between activation and task performance were observed in bilateral pre-supplementary motor area and left middle frontal gyrus. Within the dlPFC, HCP participants exhibited a significantly greater activation-performance relationship in bilateral middle frontal gyrus relative to QTIM Study participants.

**Conclusions:** The observation of diverging activation-performance relationships between two large datasets performing variations of the *n*-back task serves as a critical reminder for investigators to exercise caution when selecting WM tasks and interpreting neural activation in response to a WM task.

## 1. Introduction

Research into the neural correlates of human working memory (WM) in health and disease has been a major focus of functional Magnetic Resonance Imaging (fMRI) studies since the technique was pioneered in the 1990s (1–3). Deficits in WM are a hallmark of many neurological and psychiatric diseases, and understanding the neural mechanisms of these impairments is critical for the development of novel treatments and diagnostics. The field of psychiatry has thus embraced the study of WM in psychiatric patients, perhaps most notably in patients with schizophrenia. Differences between patients and control participants in blood-oxygen-level-dependent (BOLD) activation of the dorsolateral prefrontal cortex (dlPFC) or other brain regions activated by WM tasks are commonly observed, and this difference is presumed to reflect a neurobiological abnormality that has relevance to the patient deficit in task performance (4–9). However, as has been argued elsewhere (10), patient-control differences in brain imaging outcome measures do not necessarily reflect a true difference in the underlying neurobiology, and the study of WM in patients with schizophrenia has been plagued with non-replications and findings of activation differences in opposing directions in the patient sample (11–16).

One approach that can be used to more clearly demonstrate whether a difference in activation between psychiatric patients and healthy controls has neurobiological relevance to the construct being studied is to demonstrate a between-participants correlation or other association (e.g., using linear regression) between a neuroimaging dependent variable (such as dlPFC activation) and the phenomenon it is putatively related to (such as WM task performance). Indeed, while the term “neural correlate” is widely used in the literature to refer to a change in activation in the brain under task load (e.g., dlPFC activation is a neural correlate of WM), an empirical demonstration of such an effect does not imply an actual *correlation* between the two phenomena under study (i.e., activation and task performance). Perhaps surprisingly, within the WM literature, reports of individual-differences correlations or associations between brain activation measured with task fMRI and performance on the WM task itself are relatively uncommon. After nearly three decades and hundreds of studies, only a small handful appear have reported an association between greater activation of dlPFC and WM task performance (17–23), with a similar number reporting that the strength of deactivation of medial prefrontal cortex (mPFC) is also associated with better WM task performance (14, 15, 19, 24–26), although only the latter finding seems to have been demonstrated in patients with schizophrenia. This is arguably a “missing link” in the literature that would permit a clearer interpretation of the functional significance of patient-control differences in task activation: In the absence of a well-characterized brain-behavior relationship between activation of a brain region and performance on the task, it is difficult to know whether any abnormalities in brain activation that are observed in a patient population have relevance to their cognitive deficits.

In healthy participants, early (and subsequent) work identified a network of brain regions that is consistently activated during the performance of WM tasks, predominantly made up of regions from what is now known as the frontoparietal control network (27, 28), including bilateral dlPFC, pre-supplementary motor area (pre-SMA, sometimes referred to as dorsal anterior cingulate cortex), and intraparietal sulcus (IPS), with activation also observed in regions of the dorsal attention network (supplementary motor cortex and superior parietal lobe), cingulo-opercular network (anterior insula and additional regions of pre-SMA), and modality-specific sensory cortices (e.g., regions associated with the processing of visual or verbal stimuli).

Activation of this general network has been demonstrated with an array of different cognitive tasks that require the maintenance of information over a brief delay (ranging from ∼1-30 seconds), including the *n*-back task (27), item-recognition tasks such as the Sternberg Item Recognition Paradigm (SIRP; 28), and several other idiosyncratic WM tasks (e.g., 29, 30).

However, relatively little research has directly investigated the similarities and differences between these tasks, and in general, WM tasks are treated as relatively interchangeable, especially in the psychiatric literature investigating WM performance deficits in patient populations.

A particularly instructive example of this can be seen with *n*-back tasks, which, as summarized in **Figure 1**, have two major variants that share very few task properties, with the notable exception that on each trial the correct response depends on the stimulus that was presented some number (*n*) of trials previously. These two tasks appear to have been invented independently—and more than once (31). The first was developed in the 1950s in an unpublished dissertation, and then made its way into the broader literature (32–36). A version of this task later became one of the first tasks used in an fMRI study of patients with schizophrenia, at the National Institute of Mental Health (NIMH) in Daniel Weinberger’s group (3, 37, 38). We term this a *continuous delayed response n-back (NB-CDR)*, as it requires participants to *delay their response* for *n* trials, such that on each trial they make the behavioral response corresponding to the stimulus that was presented *n* trials previously (see **Figure 1**, top).

**Figure 1.**
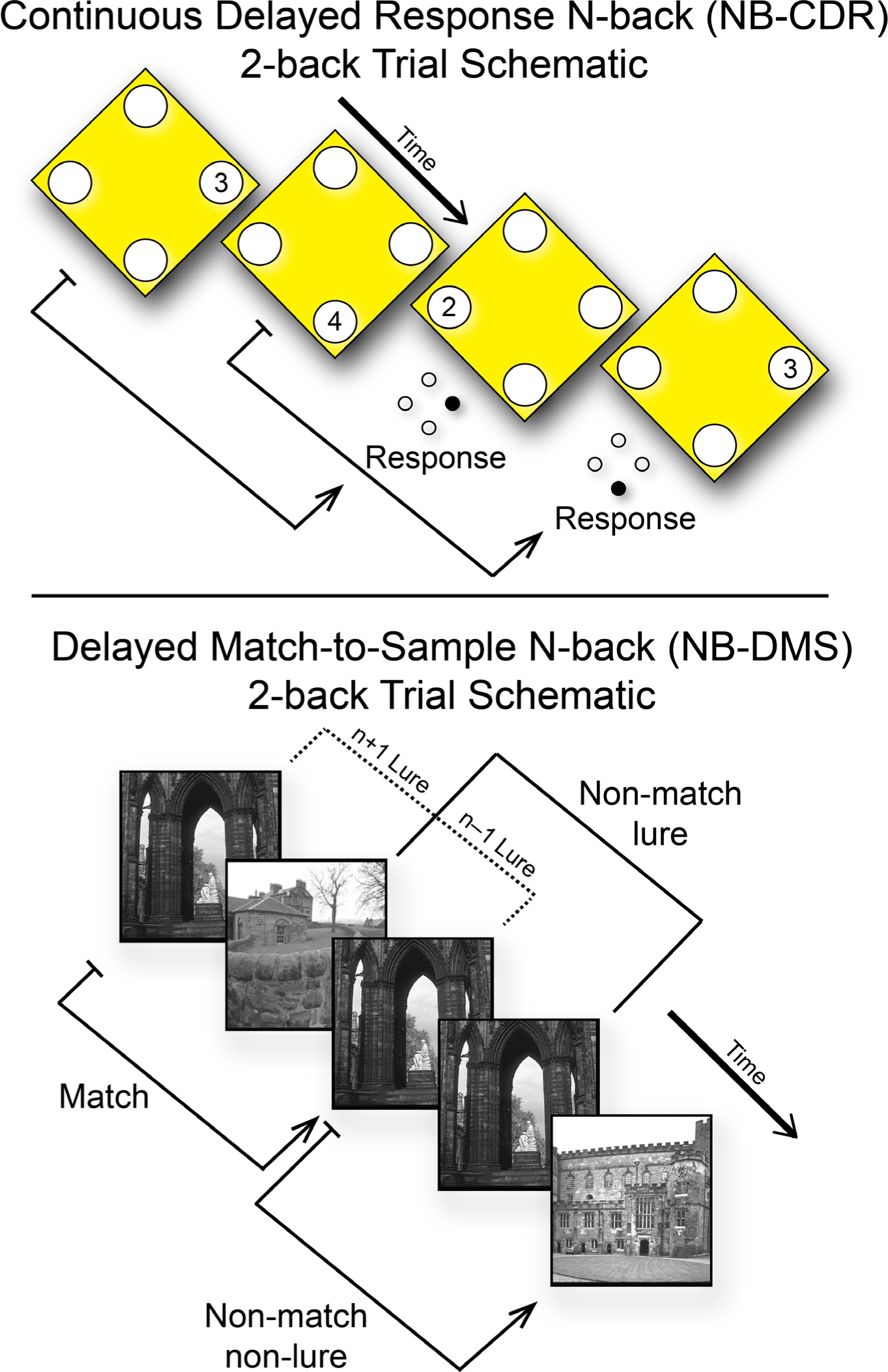
Top: Schematic of an NB-CDR 2-back trial. Stimuli typically consist of both numbers and spatial locations, and on each trial participants must make the response corresponding to the stimulus displayed n trials previously. This example reflects a sample 2-back trial of the QTIM NB-CDR task. Bottom: Schematic of an NB-DMS 2-back trial, including a recent-probe lure trial. Participants must indicate whether each presented stimulus matches the stimulus presented n trials previously. In this example, stimuli are images of places, as used in the Human Connectome Project *n*-back task.

The second, and most widely used, *n*-back was developed in the 1960s (39–41), but received relatively little attention (31). It appears to have then been independently re-invented by Alan Gevins for use in event-related potential (ERP) research in 1990 (31, 42), initiating its now widespread use in cognitive neuroscience and related fields (31, 42–48). This task involves a series of sequentially presented letters or numbers, and participants indicate whenever a stimulus matches the stimulus presented *n* trials previously; we term this a *delayed-match-to-sample n-back* (*NB-DMS*; see **Figure 1**, bottom) to distinguish it from the NB-CDR. Critically, to our knowledge, no study has ever used both tasks together, nor attempted to determine empirically which cognitive processes or brain activation patterns are shared, or distinct, between the two tasks. Notably, the correlation between dlPFC activation and WM task performance has primarily been demonstrated in NB-DMS tasks (17, 18, 20–22) and a SIRP task with distractors (19), raising the possibility that such a relationship might not exist in other WM tasks.

Consequently, we sought to utilize data from healthy participants in two large public datasets, the Human Connectome Project (HCP) 1200 Subjects Release (49–55) and the Queensland Twin Imaging (QTIM) Study (56–59), to investigate the correlation between neural activation to a WM task and in-scanner performance on that task. Furthermore, we sought to determine whether two versions of the *n*-back task, the NB-CDR and NB-DMS, elicit the same brain-behavior correlations, or whether important differences exist between these tasks in how the brain subserves task performance.

## 2. Materials and Methods

### 2.1. Participants

We utilized task-based fMRI data from 1,325 participants across two publicly available datasets: the HCP 1200 Subjects Release (n=866; 49, 50-55) and the QTIM Study (n=459; 56, 57-59). **Table 1** contains a summary of differences in demographics, data acquisition, and task design between datasets (see *Supplementary Methods* for further details).

**Table 1.** Summary of all major differences in demographics, fMRI acquisition parameters, and N-Back task design between the Human Connectome Project (HCP) and Queensland Twin Imaging (QTIM) Study datasets.

|  |  | **Dataset** |  |
| --- | --- | --- | --- |
|  |  | **HCP** | **QTIM** |
| **Demographics** | N | 866 | 459 |
| | Mean Age ($\sigma$) | 28.66 (3.74) | 22.21 (2.90) |
|  | Sex | 472 F / 394 M | 276 F / 183 M |
|  | Handedness | 785 R / 78 L / 3 NP | 459 R / 0 L |
| **fMRI Acquisition** | Scanner | Siemens Connectome Skyra 3T | Bruker MedSpec 4T |
|  | TR (ms) | 720 | 2100 |
|  | TE (ms) | 33.1 | 30 |
| | Flip Angle ($^{\circ}$) | 52 | 90 |
|  | FOV (mm) | 208 x 180 | 230 x 230 |
|  | Voxel Size (mm) | 2.0 x 2.0 x 2.0 | 3.6 x 3.6 x 3.0 |
|  | Slice Gap (mm) | 0 | 0.6 |
|  | Multiband Factor | 8 | 1 |
|  | # Volumes | 405 | 127 |
|  | Matrix Size | 104 x 90 | 64 x 64 |
| **Task Design** | # Runs | 2 | 1 |
|  | Run Length (min:sec) | 4:52 | 4:27 |
|  | Task Conditions | 0-back, 2-back | 0-back, 2-back |
|  | # Task Block per Run | 8 (4 blocks/condition) | 16 (8 blocks/condition) |
|  | # Trials per Block | 10 | 16 |
|  | Block Duration (s) | 25 | 16 |
|  | Trial Duration (s) | 2.5 | 1 |
|  | Stimulus Presentation Time (s) | 2 | 0.2 |
|  | Interstimulus Interval (s) | 0.5 | 0.8 |
|  | Stimulus Pool | Faces, Tools, Places, Body Parts | Numbers 1-4 in diamond configuration |
|  | # Potential Responses | 2 | 4 |
|  | Pre-Block Cue Duration (s) | 2.5 | 0 |
|  | Fixation Blocks? | 4/run (15 s each) | None |

HCP = Human Connectome Project; QTIM = Queensland Twin Imaging; TR = Repetition Time; TE = Echo Time; FOV = Field of View

### 2.2. Task Procedures

Participants in both datasets performed variations of a *n*-back task with different stimuli, timing, and response paradigms. Major differences between tasks are described in the *Task Design* section of **Table 1**, and examples of task response paradigms are shown in **Figure 1**. In the QTIM Study NB-CDR task, participants were presented with random sequences of the numbers 1-4 arranged in a diamond formation and responded using a response box with four buttons in the same diamond formation (**Figure 1**, top; 58, 59). For the 0-back condition, participants were asked to respond with the currently presented stimulus, while for the 2-back condition participants responded with the stimulus that was presented two stimuli prior.

In the HCP NB-DMS task (50), stimuli consisted of individually presented pictures of faces, places, tools, or body parts. In the 0-back condition, participants were presented with a ‘target’ stimulus in the beginning of the block; they were asked to respond ‘target’ on a two-button response device to any subsequent presentation of that stimulus, and ‘non-target’ everywhere else. In the 2-back condition, participants responded ‘target’ when the current stimulus matched the stimulus presented two stimuli prior (**Figure 1**, bottom).

For both tasks, performance was assessed via proportion correct and median reaction time (RT) for each task condition and across both conditions.

### 2.3. fMRI Procedures

Acquisition parameters for each dataset are detailed in **Table 1** and described in detail elsewhere (50, 59, 60). There are several major acquisition differences between the two datasets. Notably, HCP data was collected using simultaneous multi-slice (multiband) acquisition (61, 62), while QTIM Study data was collected using single-slice (single-band) acquisition. The use of multiband acceleration in HCP allowed for a shorter TR and smaller voxels without slice gaps. Another notable difference is the disparity in data quantity, where HCP participants performed two task runs, equating to over twice the duration and six times the number of volumes acquired for each QTIM Study participant.

### 2.4. fMRI Preprocessing

HCP data was obtained in its preprocessed form (55) and was preprocessed using the HCP Minimal Preprocessing Pipelines, as described elsewhere (54). Structural MRI and fMRI data from the QTIM Study were preprocessed using fMRIPrep 22.0.1 (see *Supplementary Methods*; 63, 64, 65).

### 2.5. Within-Participant Modeling

Within-participant modeling in the HCP dataset was performed as described elsewhere (50). Cortical [2-back − 0-back] contrast maps (*x[subjectID]_tfmri_wm_level2_2bk_0bk_hp200_s4_msmall*) were extracted in CIFTI format. The QTIM Study NB-CDR task was modeled using a block design with separate regressors of interest for 0-back trials, 2-back trials, and motor responses, as well as nuisance regressors (see *Supplementary Methods*). Regressors of interest were convolved with a three-parameter hemodynamic response function (HRF) with time and dispersion derivatives, as in prior work (14, 66), while nuisance regressors were not. Activation at each cortical greyordinate was quantified as the total area under the curve of the three-parameter HRF with a 2-9 second window (66, 67). Cortical [2-back − 0-back] contrasts were then calculated for each participant.

### 2.6. Regions of Interest

ROI masks of bilateral dlPFC and mPFC were identified using the parcellations (*Gordon333.32k_fs_LR*) developed by Gordon and associates (68) derived from resting-state correlations (see *Supplementary Methods,* as well as prior work (26) for details).

### 2.7. Between-Participants Analysis Within Datasets

Within each dataset, across-participant analyses of [2-back − 0-back] contrasts were performed using Permutation Analysis of Linear Models (PALM) alpha119 (RRID:SCR_017029; 69, 70) with 20,000 permutations or sign-flips using familywise error rate (FWER) correction(71) and threshold-free cluster enhancement (TFCE; 72). Exchangeability blocks were used in the HCP dataset (see *Supplementary Methods*). All resulting t-statistic maps were thresholded (FWER-corrected p < 0.05 over contrasts (73) and Dunn–Šidák corrected (74) across hemispheres).

The first analysis identified regions of significant [2-back − 0-back] activation and deactivation with a design matrix containing only an intercept. This analysis was performed across the entire cortical surface.

The second analysis determined regions with significant associations between task performance, measured as 2-back proportion correct, and [2-back − 0-back] activation, using a design matrix containing predictors for performance, age, gender, and performance*age and performance*gender interactions. All predictors were mean-centered prior to the calculation of interaction terms. This analysis was performed separately within the dlPFC and mPFC masks (see above, *2.6. Regions of Interest*), and across the entire cortical surface.

For characterization of significant greyordinates, the peak statistically significant t-statistic within each cortical parcel in the Cole-Anticevic Brain Network Parcellation (CAB-NP; 75, 76), as well as the percentage of significant greyordinates within each cortical parcel, were calculated.

### 2.8. Between-Participants Analysis Across Datasets

A between-dataset analysis of [2-back − 0-back] contrasts was also performed to identify differences in performance-activation relationships that are associated with which variation of the *n*-back task was performed. This was achieved using PALM using identical methods as described previously (see above, *2.7. Between-Participants Analysis Within Datasets*). The design matrix contained predictors for dataset membership, task performance, age, gender, and interaction terms for performance*age, performance*gender, dataset*performance, dataset*age, dataset*gender, dataset*performance*age, and dataset*performance*gender. The main predictor of interest was the dataset*performance interaction. Task performance was measured as 2-back proportion correct and was z-scored within each dataset. All other predictors were mean-centered prior to calculation of interaction terms. This analysis was performed within dlPFC and mPFC ROIs, and across all cortical greyordinates.

## 3. Results

### 3.1. Task Performance

Violin plots depicting proportion correct and median RT for each task condition (as well as for the overall task) in each dataset are shown in **Figure 2**. The median RTs across participants for the 0-back condition, 2-back condition, and the overall task in the QTIM Study dataset were 412.500 ms, 229.000 ms, and 359.000 ms, respectively. The median RTs across participants for the 0-back condition, 2-back condition, and the overall task in the HCP dataset were 741.906 ms, 957.656 ms, and 846.970 ms, respectively.

**Figure 2.**
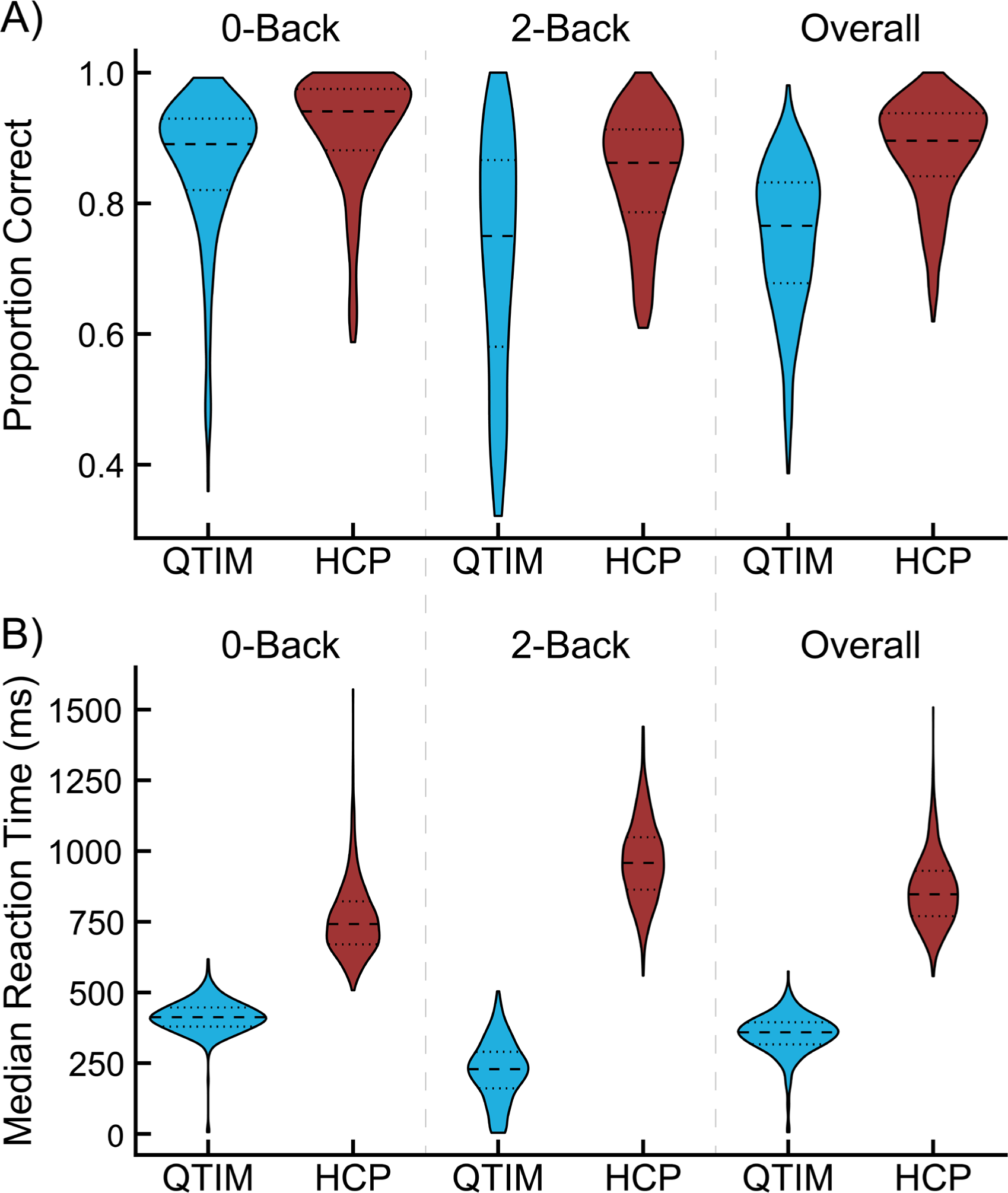
Proportion of correct responses (A) and median reaction time (B) for the 0-back condition, 2-back condition, and overall N-Back task in the Queensland Twin Imaging (QTIM) and Human Connectome Project (HCP) datasets. Central dashed lines in each violin indicate sample medians, while lower and upper dotted lines indicate 25^th^ and 75^th^ percentiles, respectively.

The median proportion of correct responses across participants for the 0-back condition, 2-back condition, and the overall task in the QTIM Study dataset were 0.891, 0.750, and 0.766, respectively. The median task accuracy across participants for the 0-back condition, 2-back condition, and the overall task in the HCP dataset were 0.941, 0.862, and 0.896, respectively.

### 3.2. Between-Participants Analysis Within Datasets

Cortical regions exhibiting significant [2-back − 0-back] activation and deactivation in each dataset are shown in **Figure 3** (peak significant t-statistics in CAB-NP parcels shown in **Figure S1** and **Figure S2**). Both datasets exhibited significant activation in bilateral dlPFC, ventrolateral PFC, pre-SMA/dorsal anterior cingulate, SMA, premotor cortex, anterior insula, IPS, precuneus, temporoparietal junction, and lateral temporal lobe. Significant deactivation was observed in bilateral mPFC, insular cortex, primary motor and somatosensory cortices, superior temporal gyrus, anterior temporal lobe and posterior cingulate cortex. Differing activation patterns were observed in primary and secondary visual cortex, with the QTIM Study dataset exhibiting deactivation in the cuneus and activation in lateral occipital lobe, and the opposite pattern in the HCP dataset.

**Figure 3.**
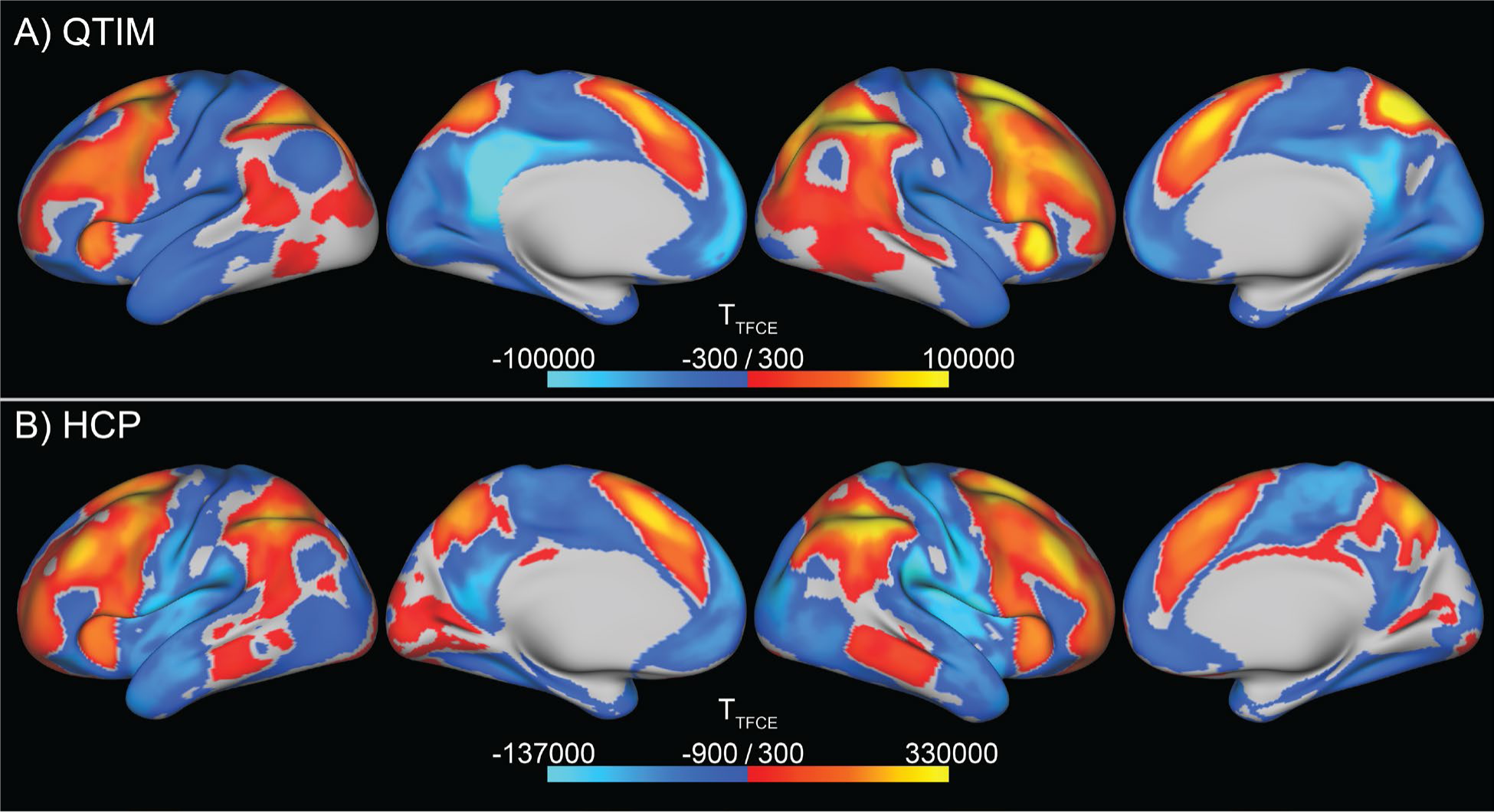
Regions across all cortical greyordinates with significant [2-back − 0-back] contrast activation (warm colors) and deactivation (cool colors) in the Queensland Twin Imaging (QTIM; A) and Human Connectome Project (HCP; B) datasets. Significant greyordinates were determined via permutation testing in Permutation Analysis of Linear Models (PALM) using 20,000 permutations, threshold-free cluster enhancement (TFCE), and family-wise error rate (FWER) correction over greyordinates and contrasts (FWER corrected, p<0.05).

Bilateral dlPFC greyordinates with significant associations between [2-back − 0-back] contrast activation and 2-back accuracy in each dataset are shown in **Figure 4** (peak significant t-statistics in CAB-NP parcels shown in **Figure S3** and **Figure S4**). The QTIM Study dataset only exhibited a positive activation-performance relationship in a small inferior region of the right precentral gyrus, and a negative relationship in the left orbitofrontal cortex (OFC) and a small region of the left middle frontal gyrus. In contrast, the HCP dataset exhibited widespread positive activation-performance relationships in a large majority of the bilateral dlPFC mask, centered on the middle frontal gyrus. A negative relationship was observed in the right ventrolateral PFC.

**Figure 4.**
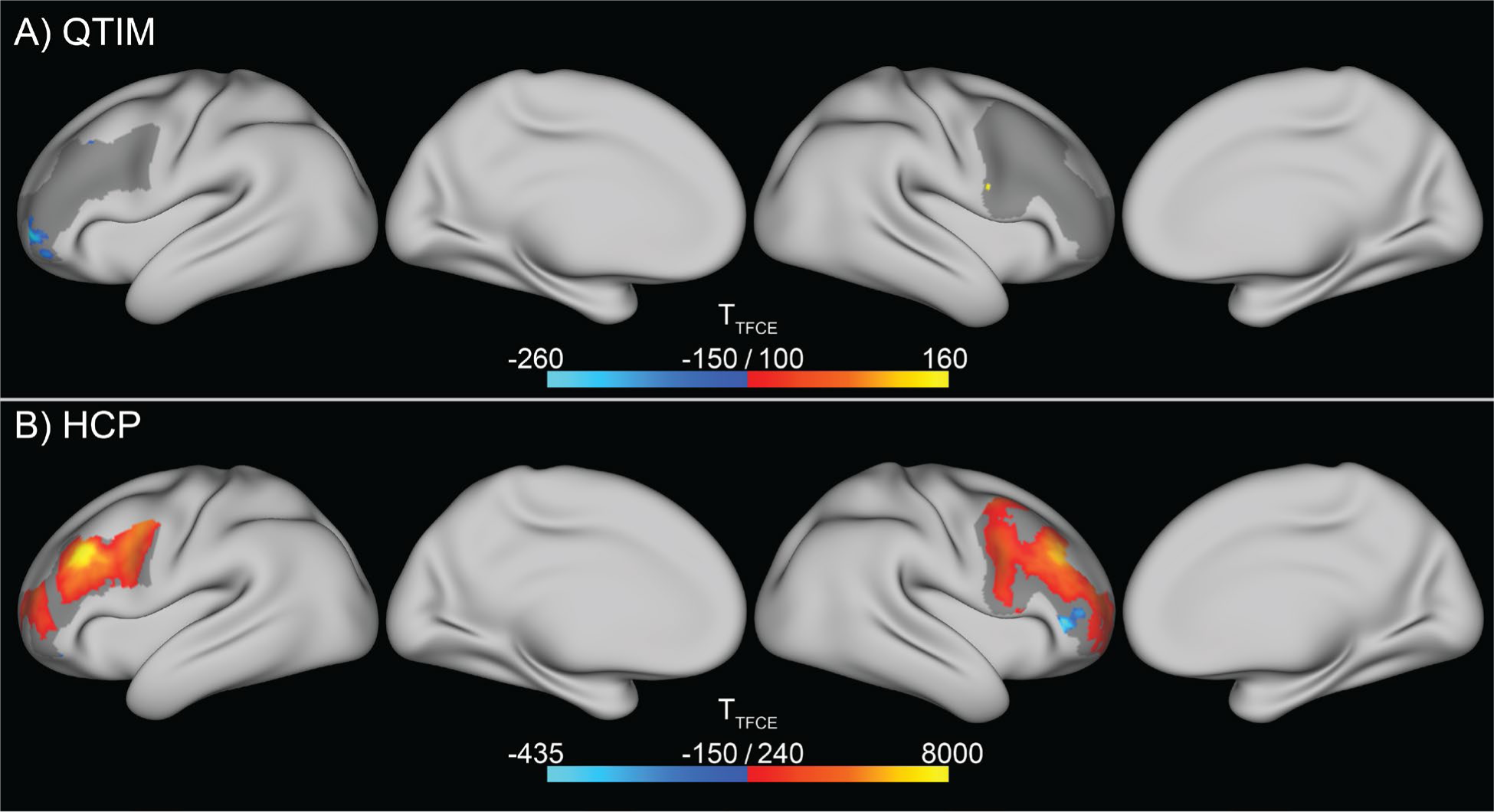
Regions with significantly positive (warm colors) and negative (cool colors) associations between [2-back − 0-back] contrast activation and 2-back accuracy in the Queensland Twin Imaging (QTIM; A) and Human Connectome Project (HCP; B) datasets. Significant greyordinates were determined within a bilateral dorsolateral prefrontal cortex (dlPFC) mask (grey underlay) via permutation testing in Permutation Analysis of Linear Models (PALM) using 20,000 permutations, threshold-free cluster enhancement (TFCE), and family-wise error rate (FWER) correction over greyordinates and contrasts (FWER corrected, p<0.05).

Greyordinates within the bilateral mPFC mask exhibiting significant activation-performance relationships are shown in **Figure 5** (peak significant t-statistics in CAB-NP parcels are shown in **Figure S5** and **Figure S6**). The QTIM Study dataset exhibited negative relationships in the anterior portion of the bilateral pre-SMA, whereas the HCP dataset showed positive relationship in this region.

**Figure 5.**
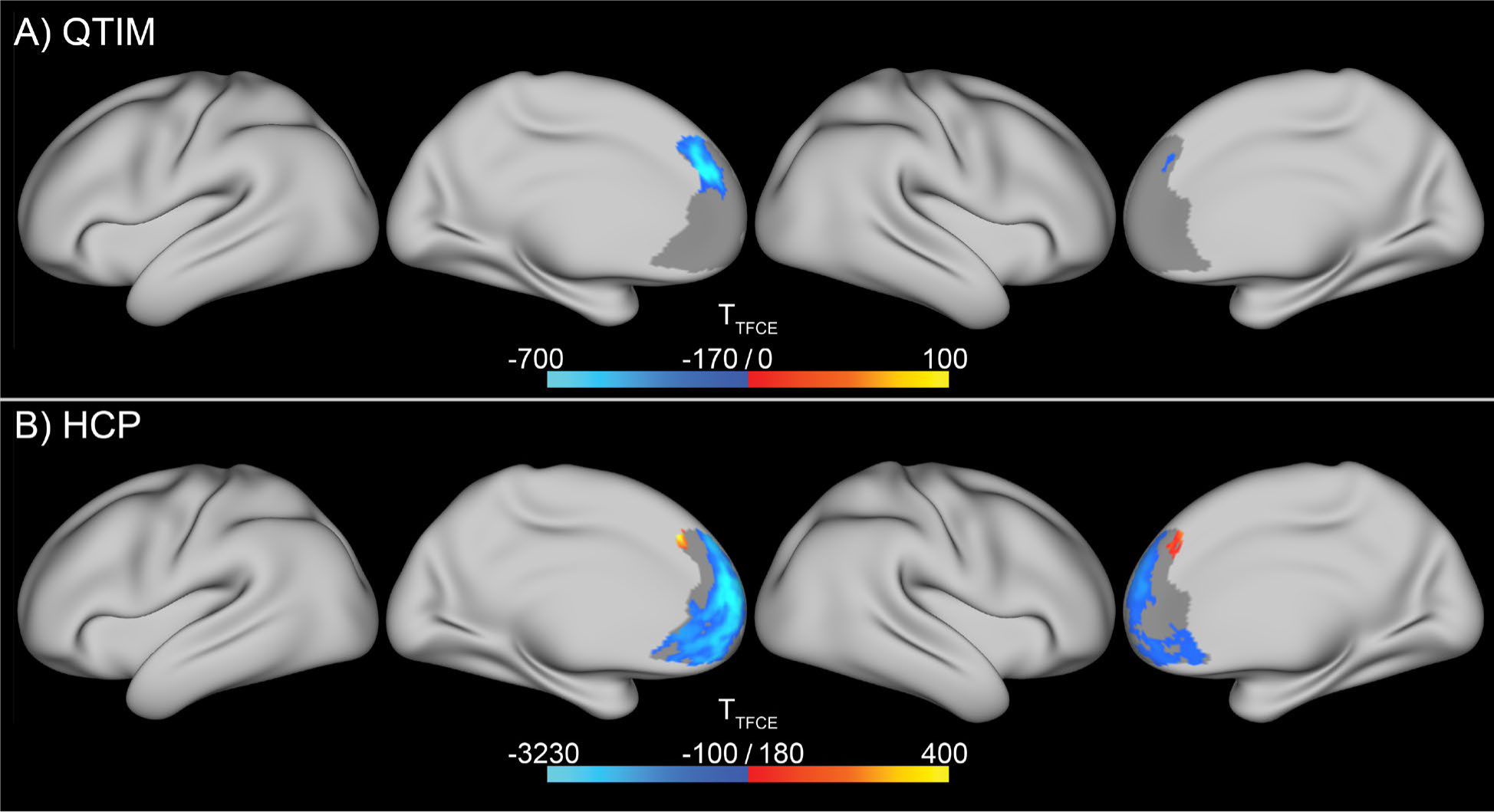
Regions with significantly positive (warm colors) and negative (cool colors) associations between [2-back − 0-back] contrast activation and 2-back accuracy in the Queensland Twin Imaging (QTIM; A) and Human Connectome Project (HCP; B) datasets. Significant greyordinates were determined within a bilateral medial prefrontal cortex (mPFC) mask (grey underlay) via permutation testing in Permutation Analysis of Linear Models (PALM) using 20,000 permutations, threshold-free cluster enhancement (TFCE), and family-wise error rate (FWER) correction over greyordinates and contrasts (FWER corrected, p<0.05). No positive associations were observed in the QTIM Study dataset.

Significant activation-performance relationships across all cortical greyordinates are shown in **Figure 6** (peak significant t-statistics in CAB-NP parcels shown in **Figure S7 and Figure S8**). The QTIM Study dataset only exhibited a positive relationship between activation and 2-back performance in a small region of the left postcentral gyrus. Negative activation-performance relationships were observed in the left middle frontal gyrus and pre-SMA. In the HCP dataset, positive activation-performance relationships were observed in bilateral dlPFC (including middle frontal gyrus), supplementary and premotor cortex, anterior poles, pre-SMA, anterior insula, middle temporal gyrus, precuneus, cuneus and calcarine fissure, right temporoparietal junction, and IPS. Negative performance-activation relationships were exhibited by mPFC, central sulcus, posterior cingulate cortex, and temporal poles.

**Figure 6.**
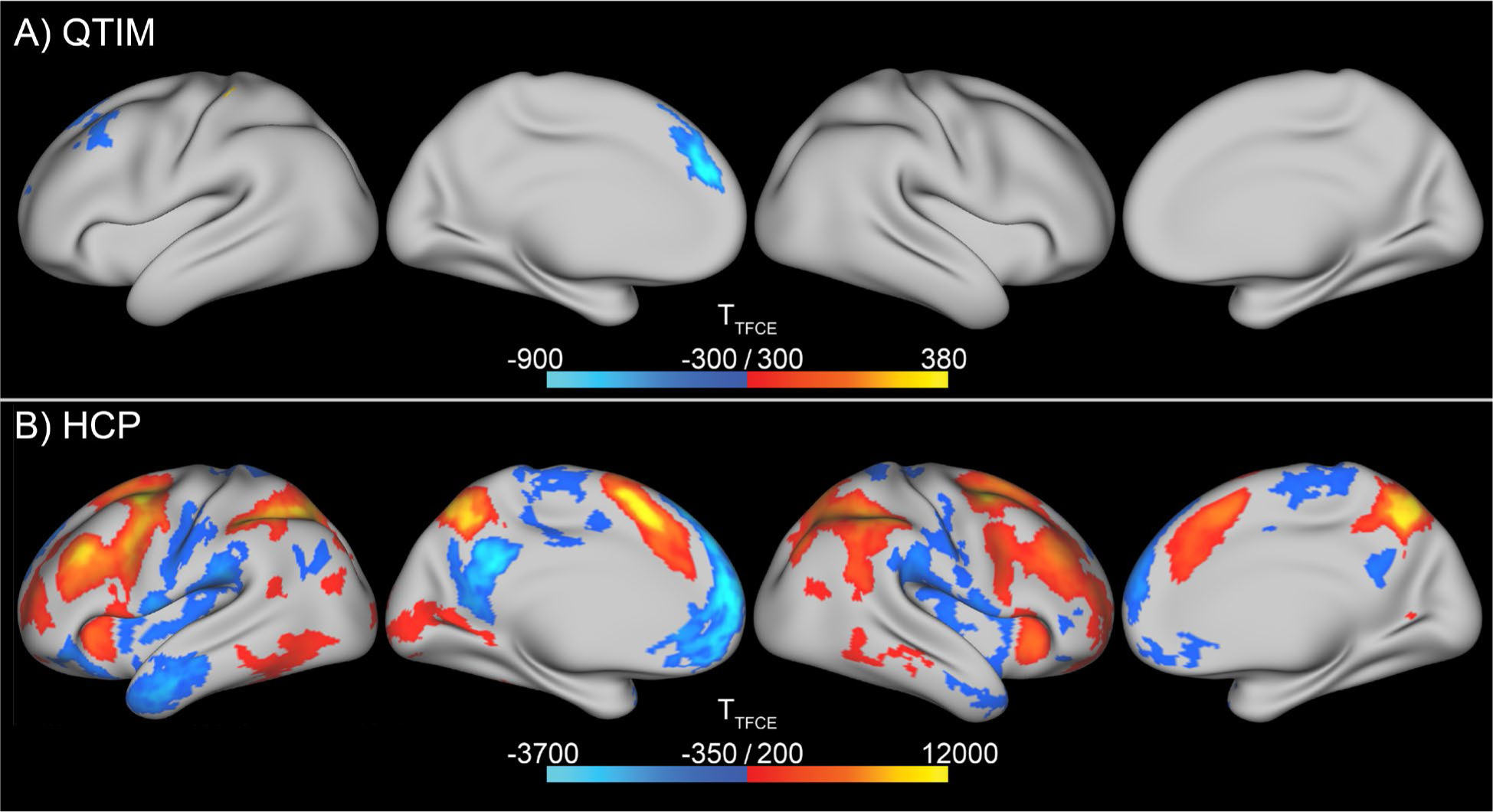
Regions across all cortical greyordinates with significantly positive (warm colors) and negative (cool colors) associations between [2-back − 0-back] contrast activation and 2-back accuracy in the Queensland Twin Imaging (QTIM; A) and Human Connectome Project (HCP; B) datasets. Results reflect a whole-brain analysis. Significant greyordinates were determined via permutation testing in Permutation Analysis of Linear Models (PALM) using 20,000 permutations, threshold-free cluster enhancement (TFCE), and family-wise error rate (FWER) correction over greyordinates and contrasts (FWER corrected, p<0.05).

### 3.3. Between-Participants Analysis Across Datasets

Cortical regions with significantly different activation-performance relationships between datasets, both within the bilateral dlPFC mask and across all cortical greyordinates, are shown in **Figure 7** (peak significant t-statistics in CAB-NP parcels shown in **Figure S9**). No significantly greater activation-performance relationships in the QTIM Study dataset over the HCP dataset were observed. Within the dlPFC mask, greater relationships in the HCP dataset over the QTIM Study dataset were observed on the bilateral middle frontal gyrus, in the location which the strongest positive activation-performance relationships were observed in the HCP dataset alone (see **Figure 4B**). Across all cortical greyordinates, greater activation-performance relationships in the HCP dataset were observed in the right precuneus and bilateral superior frontal sulcus.

**Figure 7.**
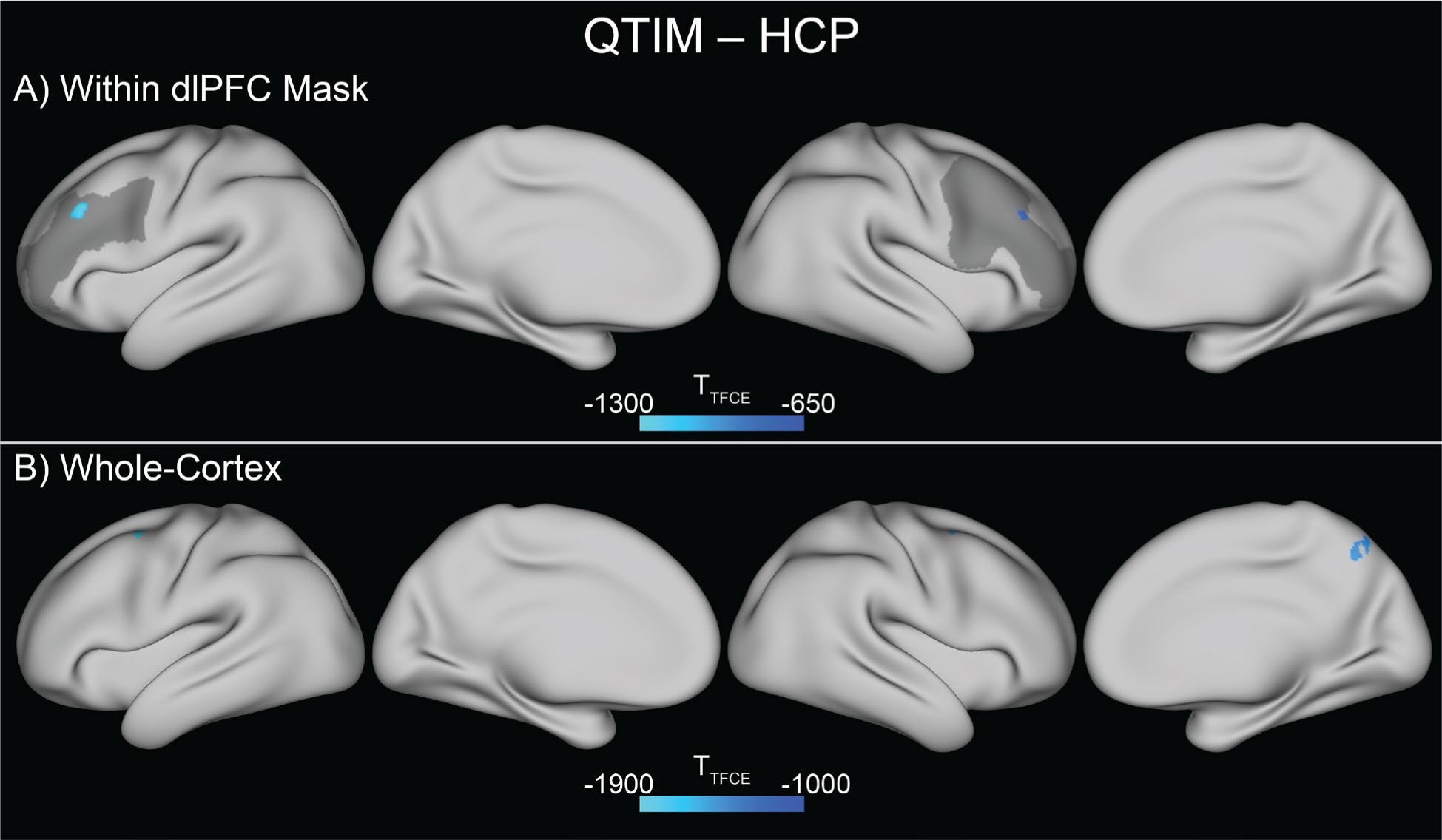
Regions with significantly lower associations between [2-back − 0-back] contrast activation and 2-back accuracy in the Queensland Twin Imaging (QTIM) dataset than the Human Connectome Project (HCP) dataset. Significant greyordinates were determined both within a bilateral dorsolateral prefrontal cortex (dlPFC) mask (A; grey underlay) and in a whole-cortex analysis (B) via permutation testing in Permutation Analysis of Linear Models (PALM) using 20,000 permutations, threshold-free cluster enhancement (TFCE), and family-wise error rate (FWER) correction over greyordinates and contrasts (FWER corrected, p<0.05). No significantly greater activation-performance associations in QTIM over HCP were identified, and thus they are not shown.

## 4. Discussion

The results presented here demonstrate a striking dissociation between BOLD estimates of neural activation and the correlation between BOLD activation and WM task performance across two versions of the *n*-back task, the NB-CDR and NB-DMS. While both tasks result in highly similar activation of the same core network of brain regions commonly observed to be active during WM task performance (along with deactivation of the default-mode network), they differ dramatically in the association between brain activation and task performance. In HCP NB-DMS task data, activation of bilateral dlPFC, frontal poles, anterior insula, pre-SMA, SMA, IPS and superior parietal lobe, middle temporal gyrus, and left visual cortex were associated with improved task performance, while deactivation of mPFC, posterior cingulate, somatomotor regions, lateral OFC, and posterior insula was also associated with improved performance.

However, these associations were largely absent in the QTIM NB-CDR task, with activation in a portion of left dlPFC and pre-SMA instead demonstrating a *negative* association with task performance. A direct comparison of these brain-behavior correlations between the two datasets revealed significant differences in bilateral dlPFC and right superior parietal lobe.

These findings highlight at least two major issues in the study of cognition and WM in both healthy and psychiatric populations. First, it is clear that the diverse array of tasks referred to as “WM tasks” in the literature should not be assumed to be interchangeable, even when they elicit activation of the same broad network of brain regions during their performance, and even when they share the same name (e.g., *n*-back). While we here characterize brain-behavior relationships in large datasets using two commonly used WM tasks, these relationships in other WM tasks (such as the commonly used SIRP) remain unknown, and could potentially even differ from the results presented here in different versions of the same tasks (e.g., letter NB-DMS tasks are extremely common in the literature, but use very different stimuli than the HCP NB-DMS used here).

Second, investigators should resist the temptation to assume that increased or decreased BOLD activation in a region that is commonly activated (or deactivated) during the performance of a cognitive task is reflective of successful task performance without directly demonstrating that that is the case. This is a particular issue in neuroimaging studies of psychiatric populations, in which increased or decreased activation by a patient population with cognitive deficits is commonly assumed to reflect a “failure to engage task networks” (in the case of decreased activation) or “inefficient cortical function” (in the case of increased activation), and such differences are broadly assumed to reflect pathology regardless of whether increased or decreased activation is observed. As demonstrated here, reduced activation in left dlPFC by a patient group could mean entirely different things depending on whether it was observed in an NB-DMS or NB-CDR task.

Unfortunately, it is impossible to pinpoint which characteristics of the NB-CDR and NB-DMS tasks, or potentially even characteristics of the participant samples or imaging acquisition procedures, led to the observed dissociation between these tasks in the relationship between BOLD activation and task performance. The HCP study was conducted on a slightly older sample of adults with a higher proportion of male participants and substantial inclusion of left-handed and ambidextrous individuals, and was undertaken using an aggressive multiband acceleration technique that led to much shorter TRs and higher spatial resolution (51, 60).

However, we view differences in the WM task used in each of the two studies as a more likely cause of the observed difference in brain-behavior correlations between the two studies, as these differences in sample and acquisition paradigm are unlikely to have produced associations with task performance in the opposite direction in some brain regions between the two datasets. In addition to major differences in the stimuli used (faces, tools, places, and body parts in the HCP study, and the numbers 1-4 combined with 4 spatial locations in the QTIM study), the QTIM study used a very rapid stimulus presentation (200 ms compared to 2000 ms in the HCP study) alongside a much shorter total trial length (1000 ms compared to 2500 ms) that could have impacted the effectiveness of any cognitive strategies used to perform the task, and thereby resulted in differences in how BOLD activation is related to task performance across individuals.

Another critical difference between the NB-CDR and NB-DMS tasks that could have led to the results reported here is that in the NB-CDR, the correct response can be prepared in advance for all task loads of 1-back or greater. An effect of this can be clearly seen in the long tails of the RT data for the NB-CDR presented in **Figure 2**, which fall below the lower limits of human RT for some participants. (We note that the long tail in RTs in the 0-back condition was driven by three participants with poor—though still above-chance—performance on that condition, and appear to have been caused by incorrect responses that were actually late responses to the preceding trial.) That is, because the NB-CDR requires participants to simply *delay* their response by *n* trials, rather than make a match/non-match discrimination on each trial based on information presented *n* trials previously, participants who have successfully encoded and maintained the stimulus can make an accurate response without even processing the stimulus presented in the current trial. When combined with fixed task presentation timing (i.e., a fixed duration and delay for each trial), participants can plan and initiate an accurate motor response before the current trial is even presented, resulting in extremely rapid RTs.

These features of the NB-CDR likely also interact with the fact that each stimulus has a 1:1 mapping to a response button, raising the possibility that participants could be preparing a sequence of motor responses that are then initiated at a fixed pace. Indeed, this set of features of the NB-CDR could be responsible for the small region of the postcentral gyrus that showed a positive association with task performance in the QTIM dataset, as well as the negative association in dlPFC and pre-SMA. That is, it may have been more optimal in the QTIM NB-CDR to use lower-order somatomotor processing to maintain prepared motor responses than it was to actively maintain a higher-order representation of prior stimuli and only initiate a response once the current stimulus is presented. Although speculative, we argue that this is the most probable explanation for the striking differences in brain-behavior associations between the NB-CDR and NB-DMS tasks observed here.

In summary, the results presented here underscore a critical consideration for the design of studies of WM, as well as for the interpretation of BOLD findings resulting from WM task-based fMRI studies of health and disease. The finding of similar patterns of task-evoked BOLD activation, yet diverging regional associations between activation and task performance, in two large datasets performing different *n*-back tasks demonstrates that investigators must exercise caution when choosing WM tasks and interpreting findings as “neural correlates” of task performance. Clearly, substantial additional work in characterizing brain-behavior relationships across the full range of WM tasks used in the literature is warranted, ideally with multiple tasks used in the same sample of participants to more clearly demonstrate which tasks or task variables (such as stimulus type or duration) result in differences in observed brain-behavior relationships, without contamination from differences in sample composition or fMRI acquisition paradigms. Additionally, we would urge researchers to systematically test for, and report, correlations between BOLD activation and task performance in their samples, in both healthy and patient samples, to develop a larger literature on these associations across the WM literature, and to provide critical context for the interpretation of patient-control differences in brain activation measures. Indeed, identifying brain regions whose activation is associated with improved performance on specific cognitive tasks could provide novel clues as to the neural basis of cognitive deficits in psychiatric patient populations with impaired cognition and disrupted activation in these regions, potentially leading to biomarker development or novel targets for pharmacological or neurostimulation interventions.

## Supporting information

Subject ID Lists

Supplementary Data from Figures

## 5. Acknowledgements

5.1. Assistance

The authors would like to acknowledge the computing resources and technical assistance provided by Stony Brook Medicine Research Computing, with substantial support from Allen Zawada and James Xikis, as well as Stony Brook Research Computing and Cyberinfrastructure and the Institute for Advanced Computational Science at Stony Brook University for access to the high-performance SeaWulf computing system (National Science Foundation Award Nos. 1531492 and 2215987, and matching funds from the Empire State Development’s Division of Science, Technology and Innovation program contract C210148), with notable support from Fırat Coşkun, Daniel Wood, and David Carlson.

## 5.3. Data Availability

Publicly available HCP data can be obtained from ConnectomeDB (https://db.humanconnectome.org/). QTIM study data can be obtained from openneuro.org (accession number *ds004169*). Subject IDs used in each dataset are available in *Supplementary Materials*.

Thresholded t-statistic maps, as well as peak significant t-statistics and percentage of significant greyordinates within each CAB-NP parcel, from all results described in Sections *3.1.* and *3.2.* are provided in CIFTI format in the *Supplementary Materials*.

## 5.4. Author Contributions

**Conceptualization:** P.N.T., J.C.W., and J.X.V.S.; **Methodology:** P.N.T., J.C.W., and J.X.V.S.; **Software:** P.N.T., J.C.W., and J.X.V.S.; **Validation:** P.N.T., J.C.W., and J.X.V.S.; **Formal Analysis:** P.N.T.; **Investigation:** P.N.T., J.C.W., and J.X.V.S.; **Resources:** J.X.V.S.; **Data Curation:** P.N.T. and J.C.W.; **Writing – Original Draft:** P.N.T. and J.X.V.S.; **Writing – Reviewing & Editing:** P.N.T., J.C.W., and J.X.V.S.; **Supervision:** J.X.V.S.; **Project Administration:** P.N.T. and J.X.V.S.; **Funding Acquisition:** J.X.V.S. and J.C.W.

## References

1. Cohen JD, Perlstein WM, Braver TS, Nystrom LE, Noll DC, Jonides J, et al. (1997): Temporal dynamics of brain activation during a working memory task. Nature. 386:604–608.

2. Braver TS, Cohen JD, Nystrom LE, Jonides J, Smith EE, Noll DC (1997): A parametric study of prefrontal cortex involvement in human working memory. NeuroImage. 5:49–62.

3. Callicott JH, Mattay VS, Bertolino A, Finn K, Coppola R, Frank JA, et al. (1999): Physiological characteristics of capacity constraints in working memory as revealed by functional MRI. Cerebral cortex. 9:20–26.

4. Carter CS, Perlstein W, Ganguli R, Brar J, Mintun M, Cohen JD (1998): Functional hypofrontality and working memory dysfunction in schizophrenia. The American journal of psychiatry. 155:1285–1287.

5. Perlstein WM, Carter CS, Noll DC, Cohen JD (2001): Relation of prefrontal cortex dysfunction to working memory and symptoms in schizophrenia. Am J Psychiat. 158:1105–1113.

6. Perlstein WM, Dixit NK, Carter CS, Noll DC, Cohen JD (2003): Prefrontal cortex dysfunction mediates deficits in working memory and prepotent responding in schizophrenia. Biological psychiatry. 53:25–38.

7. Wu D, Jiang T (2019): Schizophrenia-related abnormalities in the triple network: A meta-analysis of working memory studies. Brain Imaging and Behavior. 14:971–980.

8. Jiang S, Yan H, Chen Q, Tian L, Lu T, Tan HY, et al. (2015): Cerebral Inefficient Activation in Schizophrenia Patients and Their Unaffected Parents during the N-Back Working Memory Task: A Family fMRI Study. PloS one. 10:e0135468.

9. Anticevic A, Repovs G, Barch DM (2013): Working memory encoding and maintenance deficits in schizophrenia: neural evidence for activation and deactivation abnormalities. Schizophrenia bulletin. 39:168–178.

10. Weinberger DR, Radulescu E (2016): Finding the Elusive Psychiatric “Lesion” With 21st-Century Neuroanatomy: A Note of Caution. American Journal of Psychiatry. 173:27–33.

11. Manoach DS, Gollub RL, Benson ES, Searl MM, Goff DC, Halpern E, et al. (2000): Schizophrenic subjects show aberrant fMRI activation of dorsolateral prefrontal cortex and basal ganglia during working memory performance. Biological psychiatry. 48:99–109.

12. Manoach DS, Press DZ, Thangaraj V, Searl MM, Goff DC, Halpern E, et al. (1999): Schizophrenic subjects activate dorsolateral prefrontal cortex during a working memory task, as measured by fMRI. Biological psychiatry. 45:1128–1137.

13. Manoach DS (2003): Prefrontal cortex dysfunction during working memory performance in schizophrenia: reconciling discrepant findings. Schizophrenia research. 60:285–298.

14. Van Snellenberg JX, Girgis RR, Horga G, van de Giessen E, Slifstein M, Ojeil N, et al. (2016): Mechanisms of Working Memory Impairment in Schizophrenia. Biological psychiatry. 80:617–626.

15. Eryilmaz H, Tanner AS, Ho NF, Nitenson AZ, Silverstein NJ, Petruzzi LJ, et al. (2016): Disrupted Working Memory Circuitry in Schizophrenia: Disentangling fMRI Markers of Core Pathology vs Other Aspects of Impaired Performance. Neuropsychopharmacology. 41:2411–2420.

16. Potkin SG, Turner JA, Brown GG, McCarthy G, Greve DN, Glover GH, et al. (2009): Working memory and DLPFC inefficiency in schizophrenia: The FBIRN study. Schizophrenia bulletin. 35:19–31.

17. Cole MW, Yarkoni T, Repovs G, Anticevic A, Braver TS (2012): Global connectivity of prefrontal cortex predicts cognitive control and intelligence. The Journal of neuroscience : the official journal of the Society for Neuroscience. 32:8988–8999.

18. Satterthwaite TD, Wolf DH, Erus G, Ruparel K, Elliott MA, Gennatas ED, et al. (2013): Functional maturation of the executive system during adolescence. The Journal of neuroscience : the official journal of the Society for Neuroscience. 33:16249–16261.

19. Wager TD, Spicer J, Insler R, Smith EE (2014): The neural bases of distracter-resistant working memory. Cognitive, affective & behavioral neuroscience. 14:90–105.

20. Lamichhane B, Westbrook A, Cole MW, Braver TS (2020): Exploring brain-behavior relationships in the N-back task. NeuroImage. 212:116683.

21. Nagel IE, Preuschhof C, Li SC, Nyberg L, Backman L, Lindenberger U, et al. (2011): Load modulation of BOLD response and connectivity predicts working memory performance in younger and older adults. J Cogn Neurosci. 23:2030–2045.

22. Hakun JG, Johnson NF (2017): Dynamic range of frontoparietal functional modulation is associated with working memory capacity limitations in older adults. Brain Cogn. 118:128–136.

23. Smucny J, Hanks TD, Lesh TA, Carter CS (2023): Altered Associations Between Task Performance and Dorsolateral Prefrontal Cortex Activation During Cognitive Control in Schizophrenia. Biol Psychiatry Cogn Neurosci Neuroimaging. 8:1050–1057.

24. Anticevic A, Repovs G, Shulman GL, Barch DM (2010): When less is more: TPJ and default network deactivation during encoding predicts working memory performance. NeuroImage. 49:2638–2648.

25. Whitfield-Gabrieli S, Thermenos HW, Milanovic S, Tsuang MT, Faraone SV, McCarley RW, et al. (2009): Hyperactivity and hyperconnectivity of the default network in schizophrenia and in first-degree relatives of persons with schizophrenia. Proceedings of the National Academy of Sciences of the United States of America. 106:1279–1284.

26. Williams JC, Zheng ZJ, Tubiolo PN, Luceno JR, Gil RB, Girgis RR, et al. (2023): Medial prefrontal cortex dysfunction mediates working memory deficits in patients with schizophrenia. Biological Psychiatry: Global Open Science. 3:990–1002.

27. Owen AM, McMillan KM, Laird AR, Bullmore E (2005): N-back working memory paradigm: a meta-analysis of normative functional neuroimaging studies. Human brain mapping. 25:46–59.

28. Rottschy C, Langner R, Dogan I, Reetz K, Laird AR, Schulz JB, et al. (2012): Modelling neural correlates of working memory: a coordinate-based meta-analysis. NeuroImage. 60:830–846.

29. Van Snellenberg JX, Slifstein M, Read C, Weber J, Thompson JL, Wager TD, et al. (2015): Dynamic shifts in brain network activation during supracapacity working memory task performance. Human brain mapping. 36:1245–1264.

30. Osaka N, Osaka M, Kondo H, Morishita M, Fukuyama H, Shibasaki H (2004): The neural basis of executive function in working memory: An fMRI study based on individual differences. NeuroImage. 21:623–631.

31. Kane M, Conway A (2016): The invention of n-back: An extremely brief history. The Winnower.

32. Kay H (1953): Experimental studies of adult learningb. Unpublished doctoral dissertation, Cambridge University.

33. Kirchner WK (1958): Age differences in short-term retention of rapidly changing information. J Exp Psychol. 55:352–358.

34. Welford AT, Cambridge. University. Psychological Laboratory. Nuffield Research Unit into Problems of Ageing. [from old catalog] (1958): Ageing and human skill. London: Published for the Trustees of the Nuffield Foundation by the Oxford University Press.

35. Singleton WT (1978): Laboratory Studies of Skill. In: Singleton WT, editor. The analysis of practical skills. Dordrecht: Springer Netherlands, pp 16–43.

36. Mackworth JF (1959): Paced memorizing in a continuous task. J Exp Psychol. 58:206–211.

37. Callicott JH, Bertolino A, Mattay VS, Langheim FJ, Duyn J, Coppola R, et al. (2000): Physiological dysfunction of the dorsolateral prefrontal cortex in schizophrenia revisited. Cerebral cortex. 10:1078–1092.

38. Callicott JH, Ramsey NF, Tallen K, Bertolino A, Knable MB, Coppola R, et al. (1998): Functional magnetic resonance imaging brain mapping in psychiatry: Methodological issues illustrated in a study of working memory in schizophrenia. Neuropsychopharmacology. 18:186–196.

39. Moore ME, Ross BM (1963): Context effects in running memory. Psychological Reports. 12:451–465.

40. Ross BM (1966): Serial order as a unique source of error in running memory. Percept Mot Skills. 23:195–209.

41. Ross BM (1966): Serial-order effects in two-channel running memory. Percept Mot Skills. 23:1099–1107.

42. Gevins AS, Bressler SL, Cutillo BA, Illes J, Miller JC, Stern J, et al. (1990): Effects of prolonged mental work on functional brain topography. Electroencephalogr Clin Neurophysiol. 76:339–350.

43. Gevins A, Cutillo B (1993): Spatiotemporal dynamics of component processes in human working memory. Electroencephalogr Clin Neurophysiol. 87:128–143.

44. Cohen JD, Forman SD, Braver TS, Casey BJ, Servan-Schreiber D, Noll DC (1994): Activation of the prefrontal cortex in a nonspatial working memory task with functional MRI. Hum Brain Mapp. 1:293–304.

45. Casey BJ, Cohen JD, Jezzard P, Turner R, Noll DC, Trainor RJ, et al. (1995): Activation of prefrontal cortex in children during a nonspatial working memory task with functional MRI. Neuroimage. 2:221–229.

46. Schumacher EH, Lauber E, Awh E, Jonides J, Smith EE, Koeppe RA (1996): PET evidence for an amodal verbal working memory system. Neuroimage. 3:79–88.

47. Jonides J, Schumacher EH, Smith EE, Lauber EJ, Awh E, Minoshima S, et al. (1997): Verbal Working Memory Load Affects Regional Brain Activation as Measured by PET. J Cogn Neurosci. 9:462–475.

48. Awh E, Jonides J, Smith EE, Schumacher EH, Koeppe RA, Katz S (1996): Dissociation of Storage and Rehearsal in Verbal Working Memory: Evidence From Positron Emission Tomography. Psychological Science. 7:25–31.

49. Van Essen DC, Smith SM, Barch DM, Behrens TEJ, Yacoub E, Ugurbil K (2013): The WU-Minn Human Connectome Project: An overview. NeuroImage. 80:62–79.

50. Barch DM, Burgess GC, Harms MP, Petersen SE, Schlaggar BL, Corbetta M, et al. (2013): Function in the human connectome: Task-fMRI and individual differences in behavior. NeuroImage. 80:169–189.

51. Uğurbil K, Xu J, Auerbach EJ, Moeller S, Vu AT, Duarte-Carvajalino JM, et al. (2013): Pushing spatial and temporal resolution for functional and diffusion MRI in the Human Connectome Project. NeuroImage. 80:80–104.

52. Glasser MF, Smith SM, Marcus DS, Andersson JL, Auerbach EJ, Behrens TE, et al. (2016): The Human Connectome Project’s neuroimaging approach. Nat Neurosci. 19:1175–1187.

53. Smith SM, Beckmann CF, Andersson J, Auerbach EJ, Bijsterbosch J, Douaud G, et al. (2013): Resting-state fMRI in the Human Connectome Project. NeuroImage. 80:144–168.

54. Glasser MF, Sotiropoulos SN, Wilson JA, Coalson TS, Fischl B, Andersson JL, et al. (2013): The minimal preprocessing pipelines for the Human Connectome Project. NeuroImage. 80:105–124.

55. Hodge MR, Horton W, Brown T, Herrick R, Olsen T, Hileman ME, et al. (2016): ConnectomeDB—Sharing human brain connectivity data. NeuroImage. 124:1102–1107.

56. Strike LT, Blokland, Gabriella A.M., Hansell, Narelle K., Martin, Nicholas G., Toga, Arthur W., Thompson, Paul M., de Zubicaray, Greig I., McMahon, Katie L., Wright, Margaret J. (2023): Queensland Twin IMaging (QTIM). OpenNeuro.

57. Strike LT, Hansell NK, Couvy-Duchesne B, Thompson PM, de Zubicaray GI, McMahon KL, et al. (2019): Genetic Complexity of Cortical Structure: Differences in Genetic and Environmental Factors Influencing Cortical Surface Area and Thickness. Cerebral Cortex. 29:952–962.

58. Blokland GAM, McMahon KL, Hoffman J, Zhu G, Meredith M, Martin NG, et al. (2008): Quantifying the heritability of task-related brain activation and performance during the N-back working memory task: A twin fMRI study. Biological Psychology. 79:70–79.

59. Blokland GAM, McMahon KL, Thompson PM, Martin NG, de Zubicaray GI, Wright MJ (2011): Heritability of Working Memory Brain Activation. Journal of Neuroscience. 31:10882–10890.

60. Ugurbil K, Xu J, Auerbach EJ, Moeller S, Vu AT, Duarte-Carvajalino JM, et al. (2013): Pushing spatial and temporal resolution for functional and diffusion MRI in the Human Connectome Project. Neuroimage. 80:80–104.

61. Moeller S, Yacoub E, Olman CA, Auerbach E, Strupp J, Harel N, et al. (2010): Multiband multislice GE-EPI at 7 tesla, with 16-fold acceleration using partial parallel imaging with application to high spatial and temporal whole-brain fMRI. Magnetic resonance in medicine : official journal of the Society of Magnetic Resonance in Medicine / Society of Magnetic Resonance in Medicine. 63:1144–1153.

62. Feinberg DA, Moeller S, Smith SM, Auerbach E, Ramanna S, Gunther M, et al. (2010): Multiplexed echo planar imaging for sub-second whole brain FMRI and fast diffusion imaging. PLoS One. 5:e15710.

63. Esteban O, Markiewicz, C. J., Goncalves, M., Provins, C., Kent, J. D., DuPre, E., Salo, T., Ciric, R., Pinsard, B., Blair, R. W., Poldrack, R. A., & Gorgolewski, K. J. (2022): fMRIPrep: a robust preprocessing pipeline for functional MRI (22.0.1). Zenodo.

64. Esteban O, Markiewicz CJ, Blair RW, Moodie CA, Isik AI, Erramuzpe A, et al. (2018): fMRIPrep: a robust preprocessing pipeline for functional MRI. Nature Methods. 16:111–116.

65. Esteban O, Ciric R, Finc K, Blair RW, Markiewicz CJ, Moodie CA, et al. (2020): Analysis of task-based functional MRI data preprocessed with fMRIPrep. Nat Protoc. 15:2186–2202.

66. Van Snellenberg JX, Slifstein M, Read C, Weber J, Thompson JL, Wager TD, et al. (2015): Dynamic shifts in brain network activation during supracapacity working memory task performance. Human brain mapping. 36:1245–1264.

67. Van Snellenberg JX, Slifstein M, Read C, Weber J, Thompson JL, Wager TD, et al. (2014): Dynamic shifts in brain network activation during supracapacity working memory task performance. Human Brain Mapping. 36:1245–1264.

68. Gordon EM, Laumann TO, Adeyemo B, Huckins JF, Kelley WM, Petersen SE (2016): Generation and Evaluation of a Cortical Area Parcellation from Resting-State Correlations. Cerebral Cortex. 26:288–303.

69. Winkler AM, Ridgway GR, Webster MA, Smith SM, Nichols TE (2014): Permutation inference for the general linear model. NeuroImage. 92:381–397.

70. Winkler AM, Webster MA, Brooks JC, Tracey I, Smith SM, Nichols TE (2016): Non-parametric combination and related permutation tests for neuroimaging. Human Brain Mapping. 37:1486–1511.

71. Holmes AP, Blair RC, Watson JDG, Ford I (2016): Nonparametric Analysis of Statistic Images from Functional Mapping Experiments. Journal of Cerebral Blood Flow & Metabolism. 16:7–22.

72. Smith S, Nichols T (2009): Threshold-free cluster enhancement: Addressing problems of smoothing, threshold dependence and localisation in cluster inference. NeuroImage. 44:83–98.

73. Alberton BAV, Nichols TE, Gamba HR, Winkler AM (2020): Multiple testing correction over contrasts for brain imaging. NeuroImage. 216.

74. Šidák Z (1967): Rectangular Confidence Regions for the Means of Multivariate Normal Distributions. Journal of the American Statistical Association. 62:626–633.

75. Ji JL, Spronk M, Kulkarni K, Repovš G, Anticevic A, Cole MW (2019): Mapping the human brain’s cortical-subcortical functional network organization. NeuroImage. 185:35–57.

76. Glasser MF, Coalson TS, Robinson EC, Hacker CD, Harwell J, Yacoub E, et al. (2016): A multi-modal parcellation of human cerebral cortex. Nature. 536:171–178.

