## Supplemental Information for "A tale of two n-backs: Diverging associations of dorsolateral prefrontal cortex activation with n-back task performance"

Tubiolo *et al.*

**These supplementary materials include:**

Supplementary Methods

Supplementary Figures S1 – S9

Supplementary References

### Supplementary Methods

#### *Participants*

##### *Queensland Twin Imaging (QTIM) Study*

Unprocessed structural MRI and task-based fMRI data from 592 unrelated participants of at least 18 years of age from the QTIM Study (1-4) was obtained from *openneuro.org(RRID:SCR_005031; accession number ds004169; 5)*. Participants performed a NB-CDR task described in detail below (see main text, *2.2. Task Procedures*). From this original sample, an additional 133 participants were excluded due to poor data quality (as determined by rigorous visual quality control evaluations following preprocessing; see main text, *2.4. fMRI Preprocessing*) or poor task performance. Task performance thresholds were determined as the proportion of correct trials on each task condition needed for each individual subject to score above chance at 95% confidence according to a binomial cumulative distribution function (CDF) with a probability of success of 0.25. The resulting thresholds for proportion correct were 0.3215 and 0.3214 for 0-back and 2-back conditions, respectively. These exclusion criteria yielded a final sample of 459 participants.

##### *Human Connectome Project (HCP)*

A second cohort of preprocessed structural and task-based fMRI data was obtained from the HCP 1200 Subjects Release (6-12) comprising 866 participants performing a visual NB-DMS task(7). Selected participants had 100% task completion, no QC issues A (anatomical anomalies), B (segmentation and surface QC), and C (some data acquired during periods of head coil instability), at least two resting-state runs with at least 7.5 min of data between them (this was used as a judge of quality for MSMAll registration; 13, 14), and sufficient accuracy on both task conditions as determined by a binomial CDF with a success probability of 0.5 (proportion correct of 0.587 and 0.609 for 0-back and 2-back trials, respectively). Related subjects were permitted in this sample given that complete family information (Mother ID, Father ID, and Zygosity) was present. For each participant, the *Structural Preprocessed* and *Working Memory Task fMRI Analysis – Grayordinates, 4mm Smoothing* packages were obtained from *ConnectomeDB (RRID:SCR_004830; 12)*.

#### *fMRI Preprocessing in the Queensland Twin Imaging Study*

##### *fMRIPrep Boilerplate*

Results included in this manuscript come from preprocessing performed using fMRIPrep 22.0.1(RRID:SCR_016216; 15, 16), which is based on Nipype 1.8.4 (RRID:SCR_002502; 17, 18).

##### *Anatomical data preprocessing*

A total of 1 T1-weighted (T1w) images were found within the input BIDS dataset. The T1-weighted (T1w) image was corrected for intensity non-uniformity (INU) with *N4BiasFieldCorrection* (19), distributed with ANTs 2.3.3 (RRID:SCR_004757; 20), and used as T1w-reference throughout the workflow. The T1w-reference was then skull-stripped with a Nipype implementation of the *antsBrainExtraction.sh* workflow (from ANTs), using OASIS30ANTs as target template. Brain tissue segmentation of cerebrospinal fluid (CSF), white-matter (WM) and gray-matter (GM) was performed on the brain-extracted T1w using *fast (FSL 6.0.5.1:57b01774, RRID:SCR_002823; 21)*. Brain surfaces were reconstructed using *recon-all(FreeSurfer 7.2.0, RRID:SCR_001847; 22)*, and the brain mask estimated previously was refined with a custom variation of the method to reconcile ANTs-derived and FreeSurfer-derived segmentations of the cortical gray-matter of Mindboggle(RRID:SCR_002438; 23). Volume-based spatial normalization to two standard spaces (MNI152NLin2009cAsym, MNI152NLin6Asym) was performed through nonlinear registration with *antsRegistration* (ANTs 2.3.3), using brain-extracted versions of both T1w reference and the T1w template. The following templates were selected for spatial normalization: *ICBM 152 Nonlinear Asymmetrical template version 2009c* (RRID:SCR_008796; TemplateFlow ID: MNI152NLin2009cAsym; 24), FSL's *MNI ICBM 152 non-linear 6th Generation Asymmetric Average Brain Stereotaxic Registration Model (RRID:SCR_002823; TemplateFlow ID: MNI152NLin6Asym; 25)*.

###### *Preprocessing of B0 inhomogeneity mappings*

A total of 1 fieldmap was found available within the input BIDS structure for each subject. A deformation field to correct for susceptibility distortions was estimated based on fMRIPrep's fieldmap-less approach. The deformation field is that resulting from co-registering the EPI reference to the same-subject T1w-reference with its intensity inverted (26, 27). Registration is performed with *antsRegistration* (ANTs 2.3.3), and the process regularized by constraining deformation to be nonzero only along the phase-encoding direction, and modulated with an average fieldmap template (28).

###### *Functional data preprocessing*

For each of the 2 BOLD runs found per subject (across all tasks and sessions), the following preprocessing was performed. First, a reference volume and its skull-stripped version were generated using a custom methodology of fMRIPrep. Head-motion parameters with respect to the BOLD reference (transformation matrices, and six corresponding rotation and translation parameters) are estimated before any spatiotemporal filtering using *mcflirt (FSL 6.0.5.1:57b01774; 29)*. The estimated fieldmap was then aligned with rigid-registration to the target EPI (echo-planar imaging) reference run. The field coefficients were mapped on to the reference EPI using the transform. BOLD runs were slice-time corrected to 1.02s (0.5 of slice acquisition range 0s-2.04s) using *3dTshift* from AFNI (RRID:SCR_005927; 30). The BOLD reference was then co-registered to the T1w reference using *bbregister* (FreeSurfer) which implements boundary-based registration (31). Co-registration was configured with six degrees of freedom. Several confounding time-series were calculated based on the preprocessed BOLD: framewise displacement (FD), DVARS and three region-wise global signals. FD was computed using two formulations following Power (absolute sum of relative motions; 32) and Jenkinson(relative root mean square displacement between affines; 29). FD and DVARS are calculated for each functional run, both using their implementations in Nipype (following the definitions by Power et al.; 32). The three global signals are extracted within the CSF, the WM, and the whole-brain masks. Additionally, a set of physiological regressors were extracted to allow for component-based noise correction (CompCor; 33). Principal components are estimated after high-pass filtering the preprocessed BOLD time-series (using a discrete cosine filter with 128s cut-off) for the two CompCor variants: temporal (tCompCor) and anatomical (aCompCor). tCompCor components are then calculated from the top 2% variable voxels within the brain mask. For aCompCor, three probabilistic masks (CSF, WM and combined CSF+WM) are generated in anatomical space. The implementation differs from that of Behzadi et al. in that instead of eroding the masks by 2 pixels on BOLD space, a mask of pixels that likely contain a volume fraction of GM is subtracted from the aCompCor masks. This mask is obtained by dilating a GM mask extracted from the FreeSurfer's aseg segmentation, and it ensures components are not extracted from voxels containing a minimal fraction of GM. Finally, these masks are resampled into BOLD space and binarized by thresholding at 0.99 (as in the original implementation). Components are also calculated separately within the WM and CSF masks. For each CompCor decomposition, the *k* components with the largest singular values are retained, such that the retained components' time series are sufficient to explain 50 percent of variance across the nuisance mask (CSF, WM, combined, or temporal). The remaining components are dropped from consideration. The head-motion estimates calculated in the correction step were also placed within the corresponding confounds file. The confound time series derived from head motion estimates and global signals were expanded with the inclusion of temporal derivatives and quadratic terms for each (34). Frames that exceeded a threshold of

0.5 mm FD or 1.5 standardized DVARS were annotated as motion outliers. Additional nuisance timeseries are calculated by means of principal components analysis of the signal found within a thin band (crown) of voxels around the edge of the brain, as proposed by (35). The BOLD time-series were resampled into standard space, generating a preprocessed BOLD run in *MNI152NLin2009cAsym* space. First, a reference volume and its skull-stripped version were generated using a custom methodology of fMRIPrep. The BOLD time-series were resampled onto the following surfaces (FreeSurfer reconstruction nomenclature): *fsaverage*. Grayordinates files(11) containing 91k samples were also generated using the highest-resolution *fsaverage* as intermediate standardized surface space. All resamplings can be performed with a single interpolation step by composing all the pertinent transformations (i.e. head-motion transform matrices, susceptibility distortion correction when available, and co-registrations to anatomical and output spaces). Gridded (volumetric) resamplings were performed using *antsApplyTransforms* (ANTs), configured with Lanczos interpolation to minimize the smoothing effects of other kernels (36). Non-gridded (surface) resamplings were performed using *mri_vol2surf* (FreeSurfer).

##### *Additional Preprocessing Steps for QTIM Study Data*

During fMRIPrep preprocessing of QTIM Study data, volumetric timeseries were spatially normalized to *MNI152NLin2009cAsym* template space, and left and right cortical surfaces were then extracted in *fsaverage* space. These surfaces were resampled to CIFTI format (**.dtseries.nii*; 11), with 32,492 vertices per surface, to be used for all subsequent analyses. The first five timepoints of each run in the QTIM Study dataset were removed to ensure steady-state tissue magnetization. Each surface timeseries was then normalized to mean of 100 (37) and smoothed with a 4mm full width at half-maximum Gaussian kernel. Each participant’s structural surfaces were obtained in *fsaverage* space and resampled to FreeSurfer left-right symmetric space with 32,492 vertices (*fs_LR_32k*) to match the geometry of their surface timeseries, as well as data from HCP.

##### *Regions of Interest*

Dorsolateral prefrontal cortex (dlPFC) was taken to be regions of the Gordon parcellation communities “Dorsal Attention Network” and “Frontoparietal” localized in the PFC, while the Gordon parcellation community “Default Mode Network” was limited to just the medial prefrontal cortex (mPFC). The bilateral dlPFC regions of interest (ROIs) comprised the following parcels from the Gordon (38) parcellation: 168, 240, 272, 273, 276, 277, 319, 320, 327, 328, 236, 250, 271, 275, 74, 106, 107, 110, 113, 155, 7, 78, 108, 109, 148, 149. The bilateral mPFC ROIs comprised the following parcels: 325, 323, 322, 184, 279, 278, 25, 116, 117, 150, 152. Cortical masks of the dlPFC and mPFC were then dilated by 5 mm to fill empty space between parcels and eroded by 3 mm to remove portions of the dilated ROI falling outside the outer boundary of the chosen parcels. Dilation and erosion were performed using metric-dilate and metric-erode, respectively, in HCP Connectome Workbench v1.4.2 (RRID:SCR_008750; 39).

##### *Nuisance Regressors in Within-Participant Modeling*

In within-participant modeling in the QTIM Study dataset, nuisance regressors included six rigid-body motion parameters (along with their squares, derivatives, and squared derivatives), cerebrospinal fluid signal, and spike regressors to identify timepoints at which excessive motion occurred. Spike regressors were determined based on a run-adaptive, generalized extreme value DVARS (GEV-DV; 32, 40, 41, 42) parameter $d_G$ of 17, which flagged 2.961% of timepoints study-wide as high-motion.

##### *Exchangeability Blocks in Between-Participants Analyses*

For all between-participants analyses including participants from the HCP dataset, multi-level exchangeability blocks (43) were defined using the MATLAB (RRID:SCR_001622) code *hcp2blocks.m* provided in the PALM User Guide (https://fsl.fmrib.ox.ac.uk/fsl/fslwiki/PALM/ExchangeabilityBlocks). Blocks were defined using Mother IDs, Father IDs, and zygosity information such that families were shuffled as whole blocks, and participants within each family were only permuted amongst each other. Exchangeability blocks were not used in the QTIM Study dataset, since only unrelated participants were selected.

#### Supplementary Figures

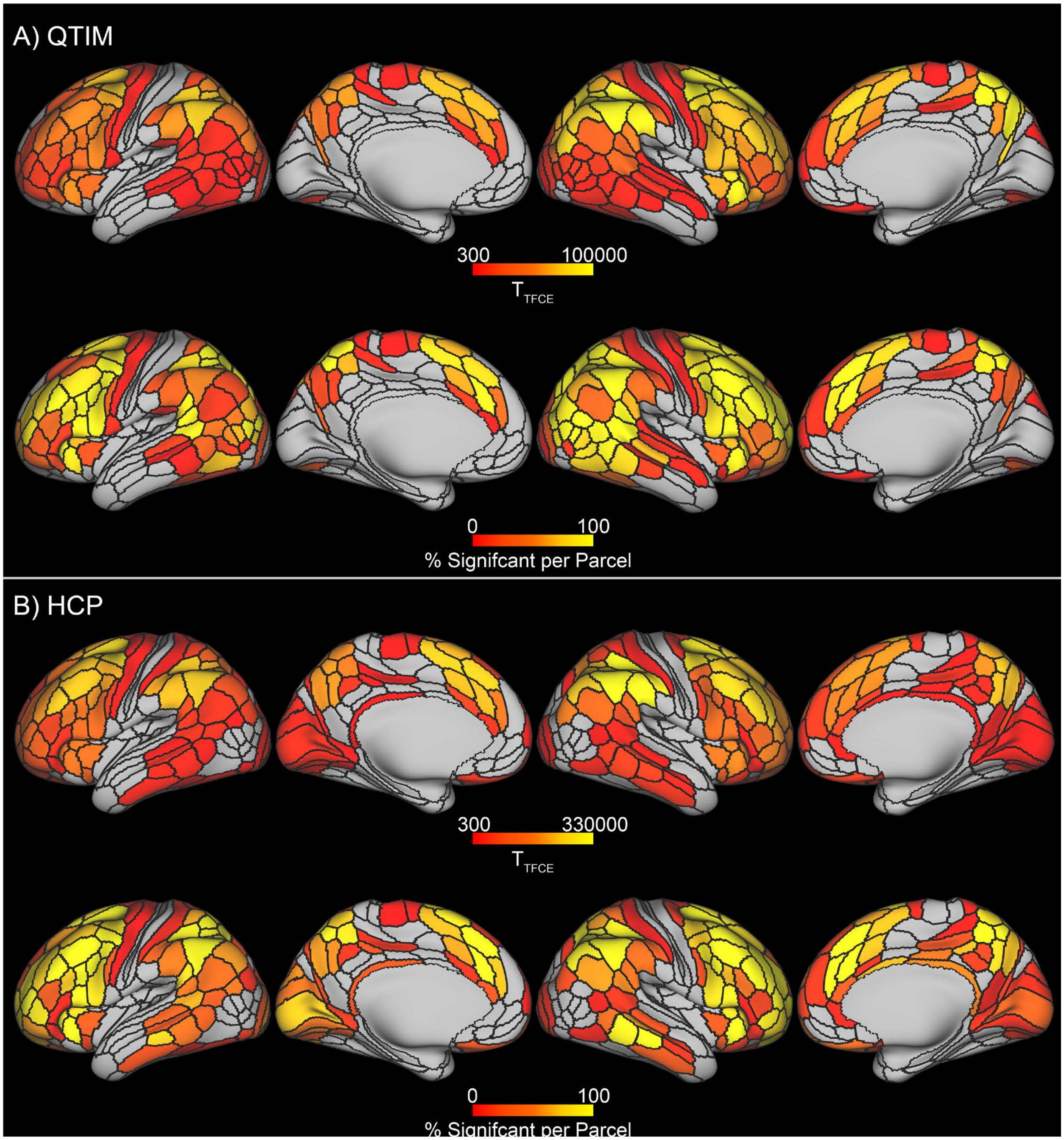


**Figure S1.** Peak t-statistic in each parcel showing significant [2-back − 0-back] contrast activation (top row of each panel), as well as the percentage of significant greyordinates within each parcel (bottom row of each panel), in the Queensland Twin Imaging (QTIM; A) and Human Connectome Project (HCP; B) datasets. Significant greyordinates were determined via permutation testing in Permutation Analysis of Linear Models (PALM) using 20,000 permutations, threshold-free cluster enhancement (TFCE), and family-wise error rate (FWER) correction over greyordinates and contrasts (FWER corrected, $p<0.05$). Resulting thresholded t-statistic maps were then parcellated using cortical regions of the Cole-Anticevic Brain Network Parcellation (CAB-NP).

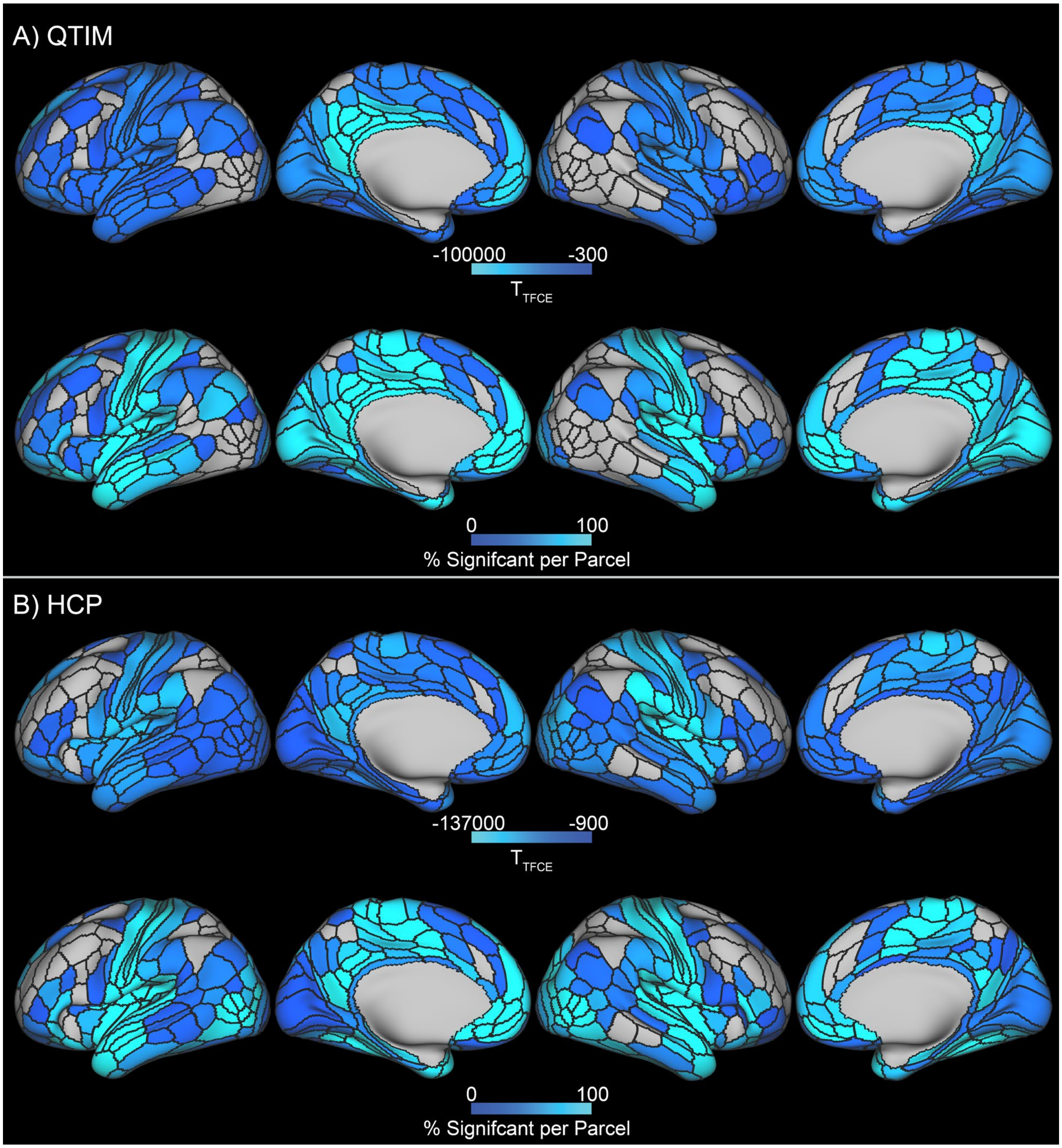


**Figure S2.** Peak t-statistic in each parcel showing significant [2-back − 0-back] contrast deactivation (top row of each panel), as well as the percentage of significant greyordinates within each parcel (bottom row of each panel), in the Queensland Twin Imaging (QTIM; A) and Human Connectome Project (HCP; B) datasets. Significant greyordinates were determined via permutation testing in Permutation Analysis of Linear Models (PALM) using 20,000 permutations, threshold-free cluster enhancement (TFCE), and family-wise error rate (FWER) correction over greyordinates and contrasts (FWER corrected, $p<0.05$). Resulting thresholded t-statistic maps were then parcellated using cortical regions of the Cole-Anticevic Brain Network Parcellation (CAB-NP).

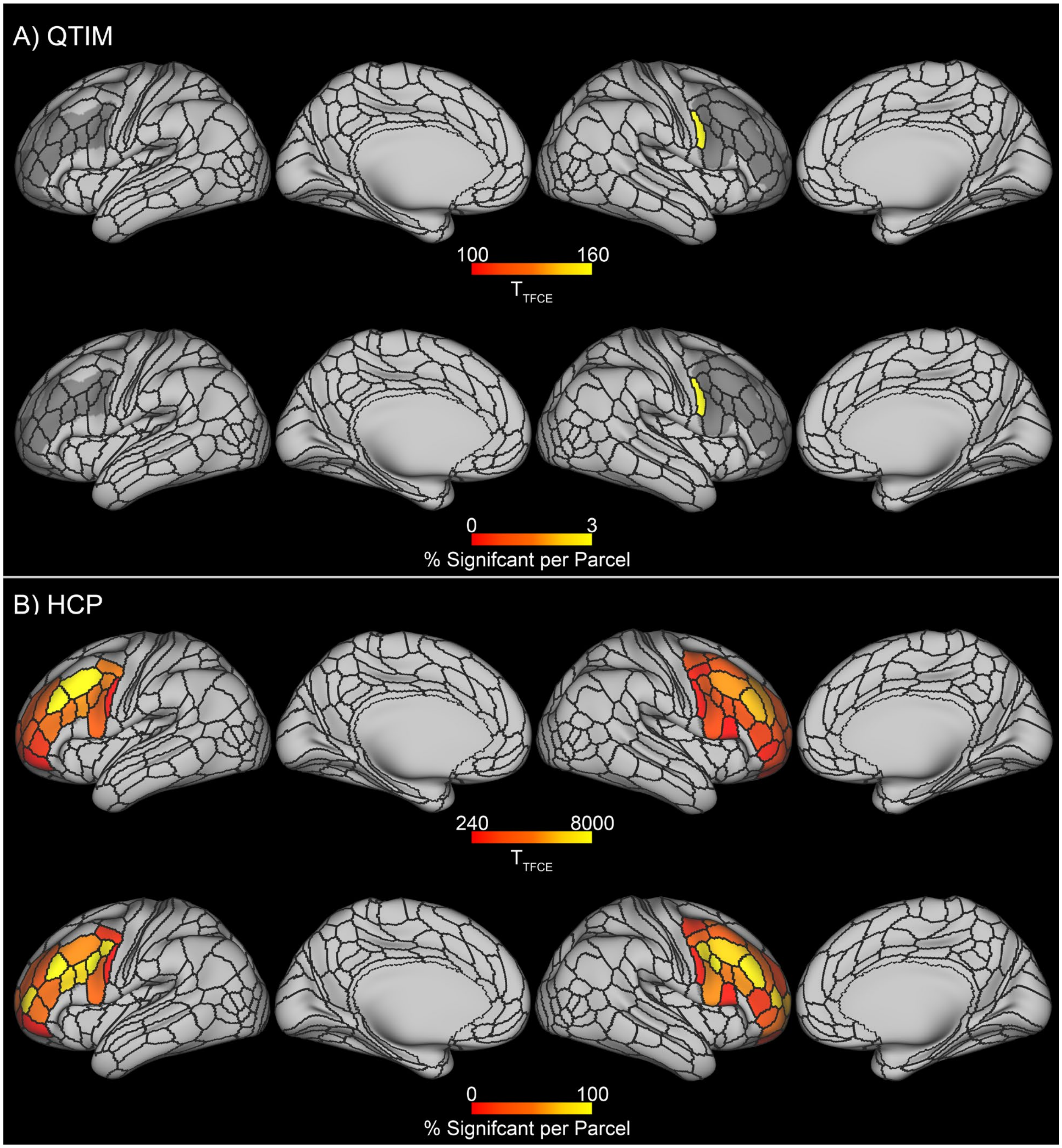


**Figure S3.** Peak t-statistic in each parcel showing significantly positive associations between [2-back − 0-back] contrast activation and 2-back accuracy (top row of each panel), as well as the percentage of significant greyordinates within each parcel (bottom row of each panel), in the Queensland Twin Imaging (QTIM; A) and Human Connectome Project (HCP; B) datasets. Significant greyordinates were determined within a bilateral dorsolateral prefrontal cortex mask (grey underlay) via permutation testing in Permutation Analysis of Linear Models (PALM) using 20,000 permutations, threshold-free cluster enhancement (TFCE), and family-wise error rate (FWER) correction over greyordinates and contrasts (FWER corrected, $p<0.05$). Resulting thresholded t-statistic maps were then parcellated using cortical regions of the Cole-Anticevic Brain Network Parcellation (CAB-NP).

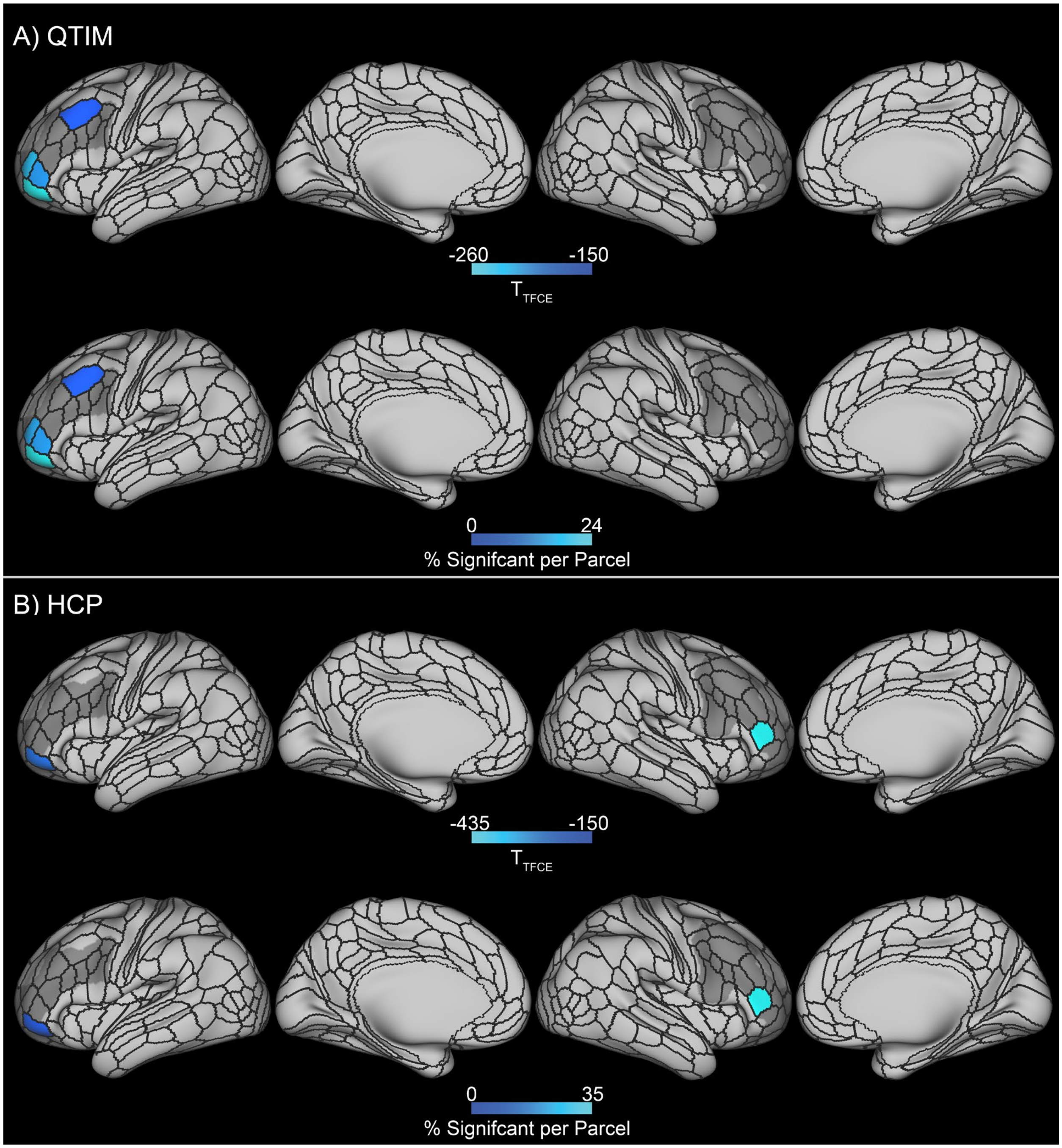


**Figure S4.** Peak t-statistic in each parcel showing significantly negative associations between [2-back − 0-back] contrast activation and 2-back accuracy (top row of each panel), as well as the percentage of significant greyordinates within each parcel (bottom row of each panel), in the Queensland Twin Imaging (QTIM; A) and Human Connectome Project (HCP; B) datasets. Significant greyordinates were determined within a bilateral dorsolateral prefrontal cortex mask (grey underlay) via permutation testing in Permutation Analysis of Linear Models (PALM) using 20,000 permutations, threshold-free cluster enhancement (TFCE), and family-wise error rate (FWER) correction over greyordinates and contrasts (FWER corrected, $p<0.05$). Resulting thresholded t-statistic maps were then parcellated using cortical regions of the Cole-Anticevic Brain Network Parcellation (CAB-NP).

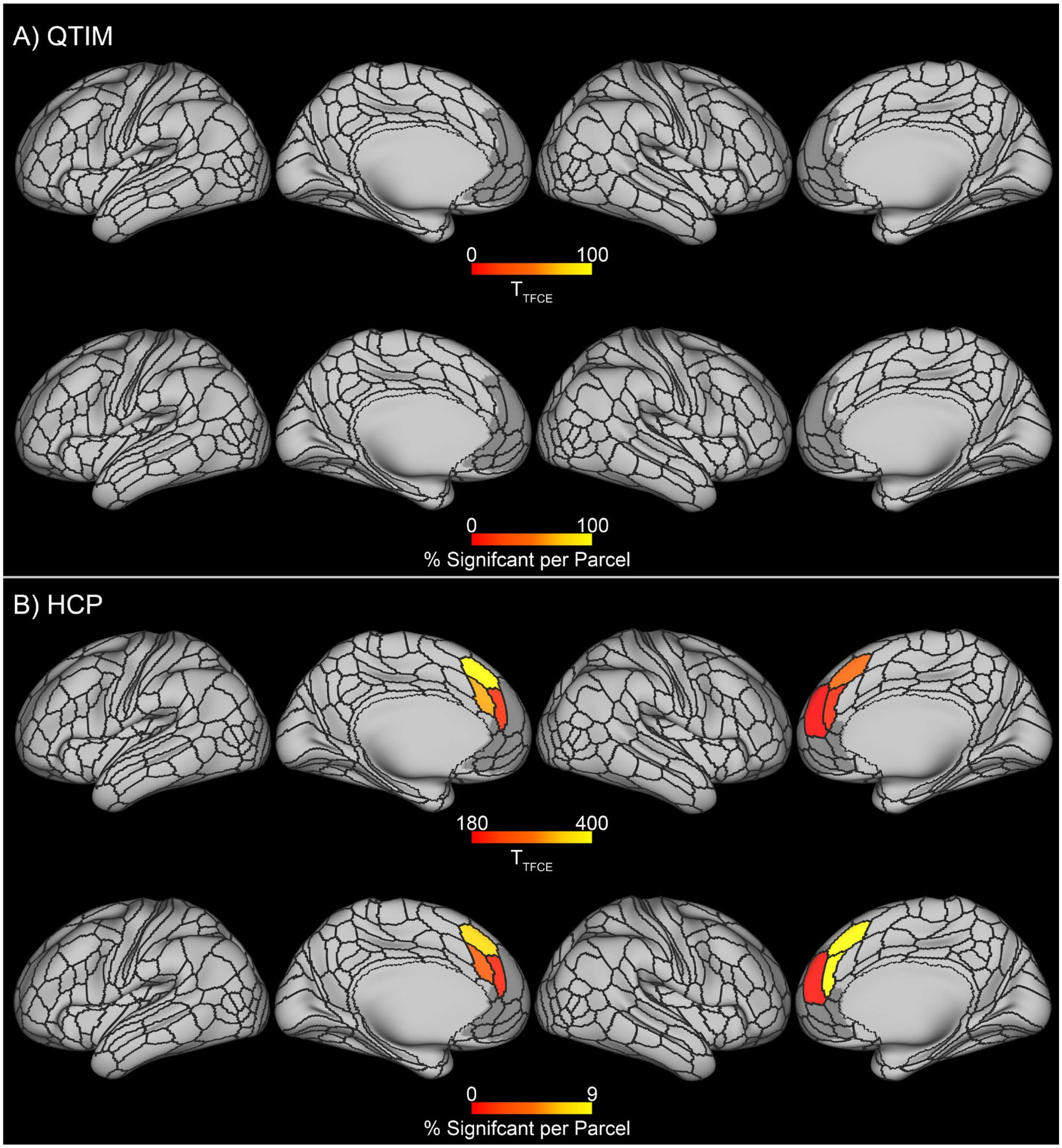


**Figure S5.** Peak t-statistic in each parcel showing significantly positive associations between [2-back − 0-back] contrast activation and 2-back accuracy (top row of each panel), as well as the percentage of significant greyordinates within each parcel (bottom row of each panel), in the Queensland Twin Imaging (QTIM; A) and Human Connectome Project (HCP; B) datasets. Significant greyordinates were determined within a bilateral medial prefrontal cortex mask (grey underlay) via permutation testing in Permutation Analysis of Linear Models (PALM) using 20,000 permutations, threshold-free cluster enhancement (TFCE), and family-wise error rate (FWER) correction over greyordinates and contrasts (FWER corrected, $p<0.05$). Resulting thresholded t-statistic maps were then parcellated using cortical regions of the Cole-Anticevic Brain Network Parcellation (CAB-NP). No significant greyordinates were found in the QTIM dataset.

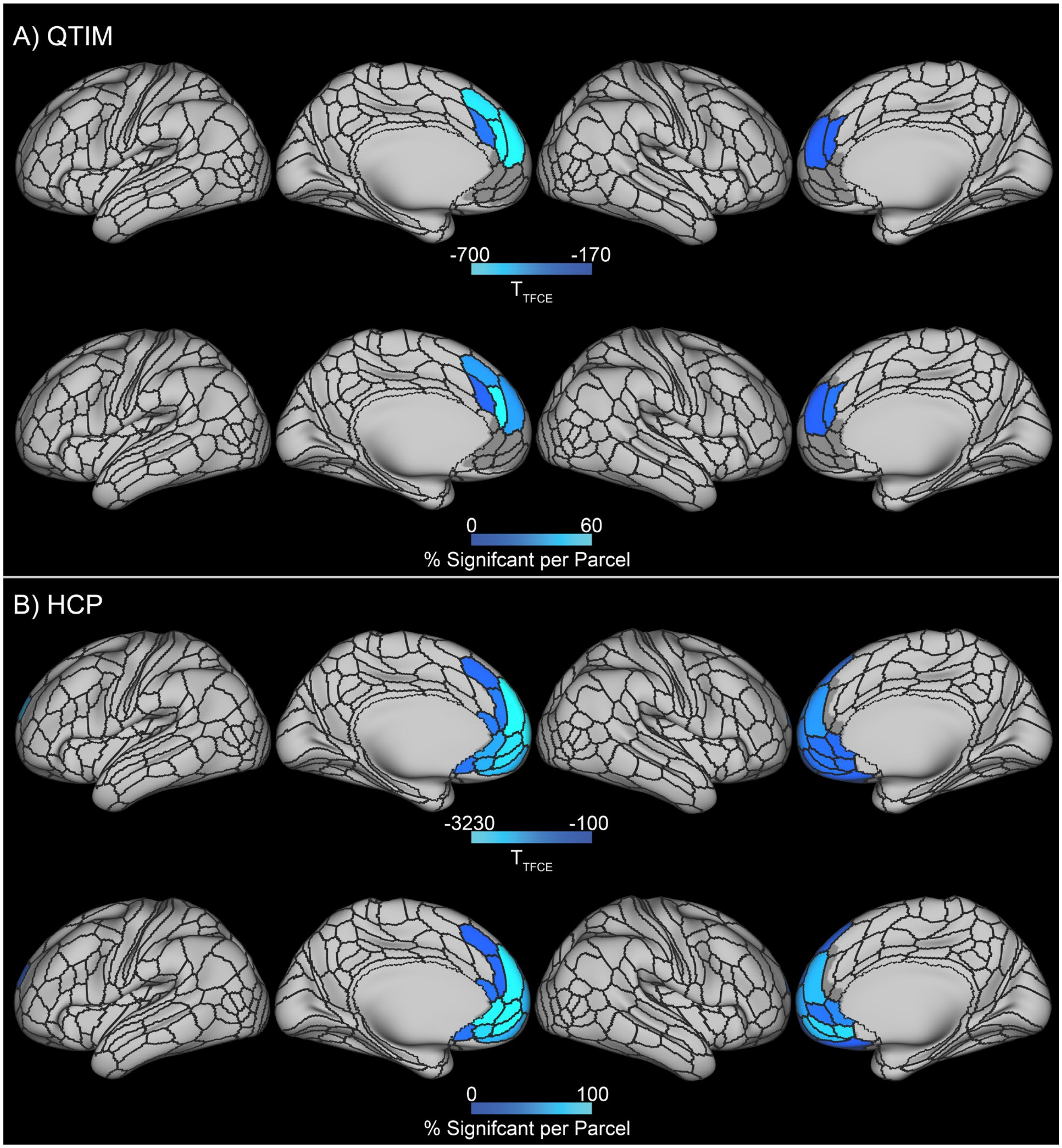


**Figure S6.** Peak t-statistic in each parcel showing significantly negative associations between [2-back − 0-back] contrast activation and 2-back accuracy (top row of each panel), as well as the percentage of significant greyordinates within each parcel (bottom row of each panel), in the Queensland Twin Imaging (QTIM; A) and Human Connectome Project (HCP; B) datasets. Significant greyordinates were determined within a bilateral medial prefrontal cortex mask (grey underlay) via permutation testing in Permutation Analysis of Linear Models (PALM) using 20,000 permutations, threshold-free cluster enhancement (TFCE), and family-wise error rate (FWER) correction over greyordinates and contrasts (FWER corrected, $p<0.05$). Resulting thresholded t-statistic maps were then parcellated using cortical regions of the Cole-Anticevic Brain Network Parcellation (CAB-NP). No significant greyordinates were found in the QTIM dataset.

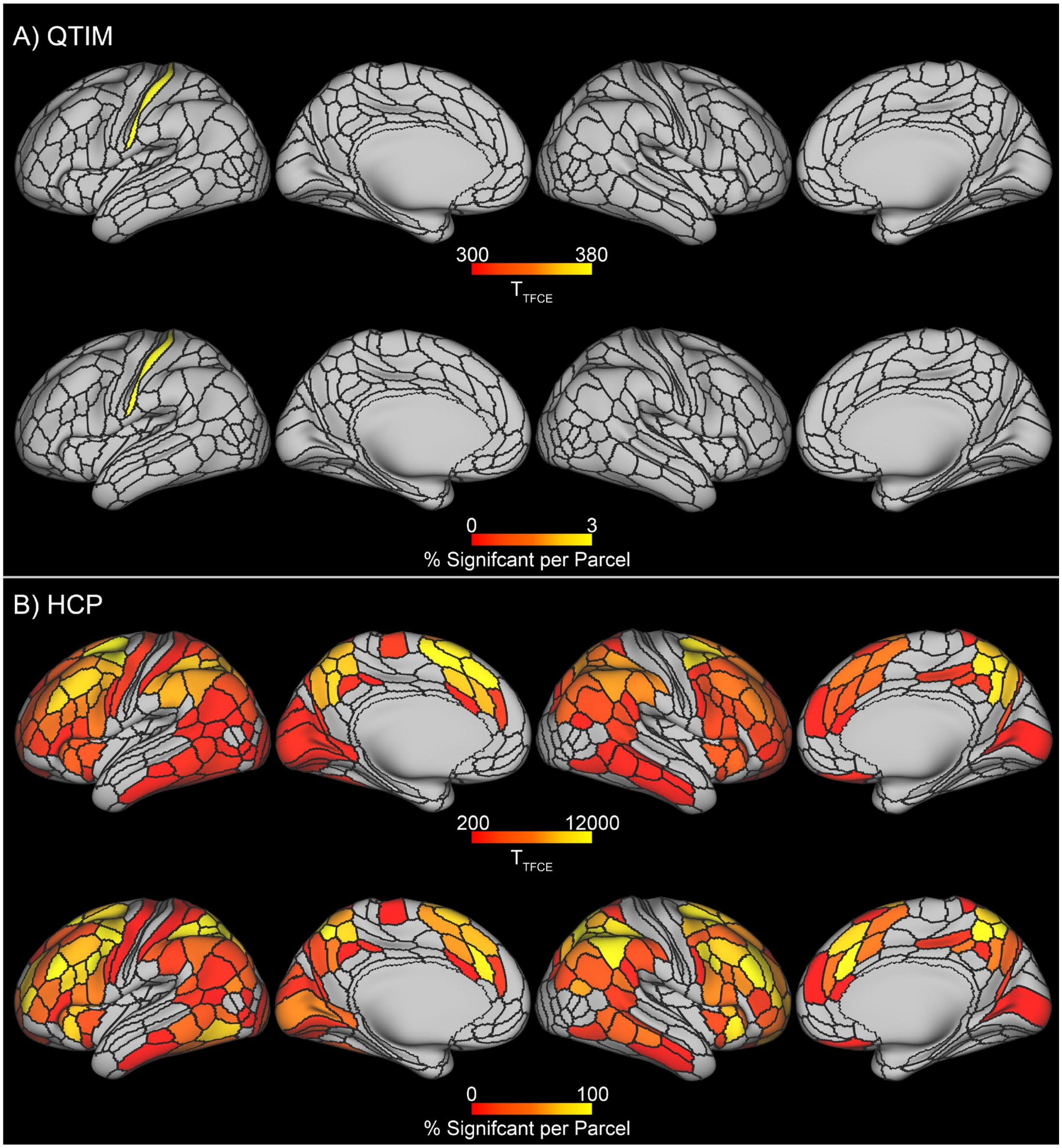


**Figure S7.** Peak t-statistic in each parcel showing significantly positive associations between [2-back − 0-back] contrast activation and 2-back accuracy (top row of each panel), as well as the percentage of significant greyordinates within each parcel (bottom row of each panel), in the Queensland Twin Imaging (QTIM; A) and Human Connectome Project (HCP; B) datasets. Significant greyordinates were determined via permutation testing in Permutation Analysis of Linear Models (PALM) using 20,000 permutations, threshold-free cluster enhancement (TFCE), and family-wise error rate (FWER) correction over greyordinates and contrasts (FWER corrected, $p<0.05$). Resulting thresholded t-statistic maps were then parcellated using cortical regions of the Cole-Anticevic Brain Network Parcellation (CAB-NP).

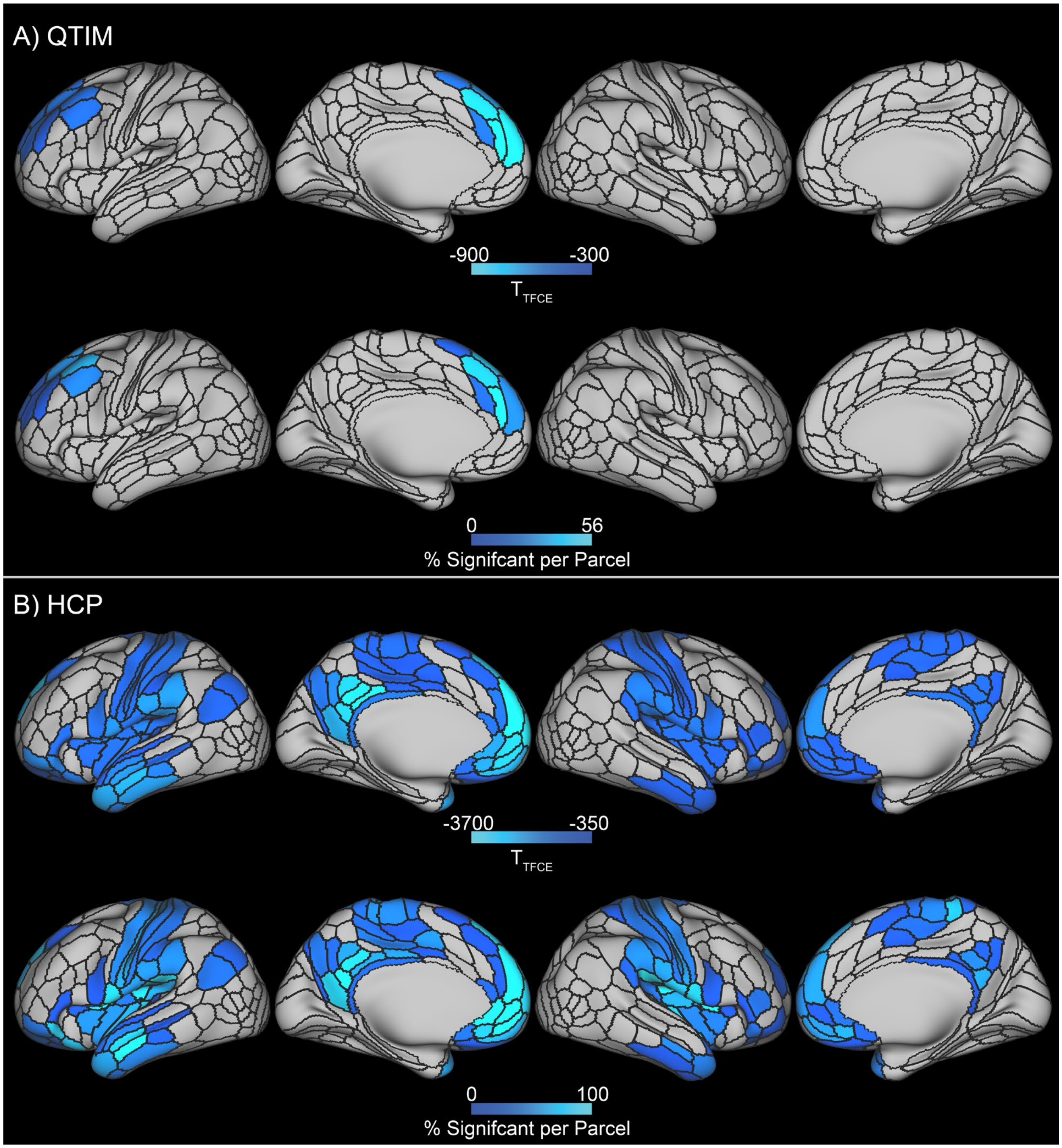


**Figure S8.** Peak t-statistic in each parcel showing significantly negative associations between [2-back − 0-back] contrast activation and 2-back accuracy (top row of each panel), as well as the percentage of significant greyordinates within each parcel (bottom row of each panel), in the Queensland Twin Imaging (QTIM; A) and Human Connectome Project (HCP; B) datasets. Significant greyordinates were determined via permutation testing in Permutation Analysis of Linear Models (PALM) using 20,000 permutations, threshold-free cluster enhancement (TFCE), and family-wise error rate (FWER) correction over greyordinates and contrasts (FWER corrected, $p<0.05$). Resulting thresholded t-statistic maps were then parcellated using cortical regions of the Cole-Anticevic Brain Network Parcellation (CAB-NP).

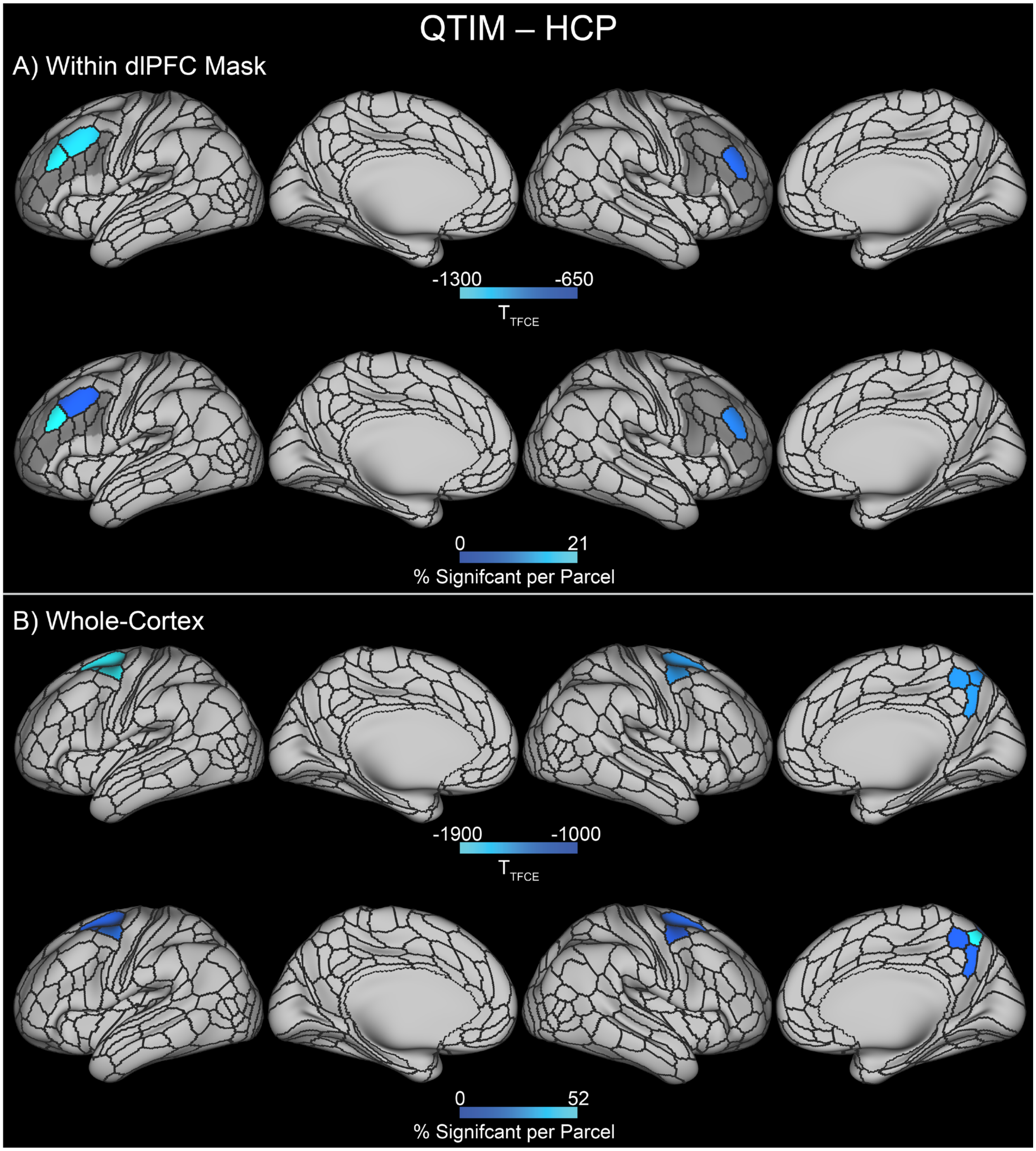


**Figure S9.** Peak t-statistic in each parcel showing significantly lower associations between [2-back − 0-back] contrast activation and 2-back accuracy in the Queensland Twin Imaging (QTIM) dataset than the Human Connectome Project (HCP) dataset (top row of each panel), as well as the percentage of significant greyordinates within each parcel (bottom row of each panel). Significant greyordinates were determined both within a bilateral dorsolateral prefrontal cortex (dlPFC) mask (A; grey underlay) and in a whole-cortex analysis (B) via permutation testing in Permutation Analysis of Linear Models (PALM) using 20,000 permutations, threshold-free cluster enhancement (TFCE), and family-wise error rate (FWER) correction over greyordinates and contrasts (FWER corrected, $p<0.05$). Resulting thresholded t-statistic maps were then parcellated using cortical regions of the Cole-Anticevic Brain Network Parcellation (CAB-NP).

#### Supplementary References


1. Strike LT, Blokland, Gabriella A.M., Hansell, Narelle K., Martin, Nicholas G., Toga, Arthur W., Thompson, Paul M., de Zubicaray, Greig I., McMahon, Katie L., Wright, Margaret J. (2023): Queensland Twin IMaging (QTIM). OpenNeuro.
2. Strike LT, Hansell NK, Couvy-Duchesne B, Thompson PM, de Zubicaray GI, McMahon KL, et al. (2019): Genetic Complexity of Cortical Structure: Differences in Genetic and Environmental Factors Influencing Cortical Surface Area and Thickness. *Cerebral Cortex*. 29:952-962.
3. Blokland GAM, McMahon KL, Hoffman J, Zhu G, Meredith M, Martin NG, et al. (2008): Quantifying the heritability of task-related brain activation and performance during the N-back working memory task: A twin fMRI study. *Biological Psychology*. 79:70-79.
4. Blokland GAM, McMahon KL, Thompson PM, Martin NG, de Zubicaray GI, Wright MJ (2011): Heritability of Working Memory Brain Activation. *Journal of Neuroscience*. 31:10882-10890.
5. Markiewicz CJ, Gorgolewski KJ, Feingold F, Blair R, Halchenko YO, Miller E, et al. (2021): The OpenNeuro resource for sharing of neuroscience data. *eLife*. 10.
6. Van Essen DC, Smith SM, Barch DM, Behrens TEJ, Yacoub E, Ugurbil K (2013): The WU-Minn Human Connectome Project: An overview. *NeuroImage*. 80:62-79.
7. Barch DM, Burgess GC, Harms MP, Petersen SE, Schlaggar BL, Corbetta M, et al. (2013): Function in the human connectome: Task-fMRI and individual differences in behavior. *NeuroImage*. 80:169-189.
8. Uğurbil K, Xu J, Auerbach EJ, Moeller S, Vu AT, Duarte-Carvajalino JM, et al. (2013): Pushing spatial and temporal resolution for functional and diffusion MRI in the Human Connectome Project. *NeuroImage*. 80:80-104.

9. Glasser MF, Smith SM, Marcus DS, Andersson JL, Auerbach EJ, Behrens TE, et al. (2016): The Human Connectome Project's neuroimaging approach. *Nat Neurosci*. 19:1175-1187.

10. Smith SM, Beckmann CF, Andersson J, Auerbach EJ, Bijsterbosch J, Douaud G, et al. (2013): Resting-state fMRI in the Human Connectome Project. *NeuroImage*. 80:144-168.

11. Glasser MF, Sotiropoulos SN, Wilson JA, Coalson TS, Fischl B, Andersson JL, et al. (2013): The minimal preprocessing pipelines for the Human Connectome Project. *NeuroImage*. 80:105-124.

12. Hodge MR, Horton W, Brown T, Herrick R, Olsen T, Hileman ME, et al. (2016): ConnectomeDB—Sharing human brain connectivity data. *NeuroImage*. 124:1102-1107.

13. Robinson EC, Garcia K, Glasser MF, Chen Z, Coalson TS, Makropoulos A, et al. (2018): Multimodal surface matching with higher-order smoothness constraints. *NeuroImage*. 167:453-465.

14. Robinson EC, Jbabdi S, Glasser MF, Andersson J, Burgess GC, Harms MP, et al. (2014): MSM: A new flexible framework for Multimodal Surface Matching. *NeuroImage*. 100:414-426.

15. Esteban O, Markiewicz, C. J., Goncalves, M., Provins, C., Kent, J. D., DuPre, E., Salo, T., Ciric, R., Pinsard, B., Blair, R. W., Poldrack, R. A., & Gorgolewski, K. J. (2022): fMRIPrep: a robust preprocessing pipeline for functional MRI (22.0.1). Zenodo.

16. Esteban O, Markiewicz CJ, Blair RW, Moodie CA, Isik AI, Erramuzpe A, et al. (2018): fMRIPrep: a robust preprocessing pipeline for functional MRI. *Nature Methods*. 16:111-116.

17. Esteban O, Markiewicz, C. J., Burns, C., Goncalves, M., Jarecka, D., Ziegler, E., Berleant, S., Ellis, D. G., Pinsard, B., Madison, C., Waskom, M., Notter, M. P., Clark, D., Manhães-Savio, A., Clark, D., Jordan, K., Dayan, M., Halchenko, Y. O., Loney, F., … Ghosh, S. (2022): nipy/nipype: 1.8.3 (1.8.3). Zenodo.

18. Ghosh SS, Waskom ML, Halchenko YO, Clark D, Madison C, Burns CD, et al. (2011): Nipype: A Flexible, Lightweight and Extensible Neuroimaging Data Processing Framework in Python. *Frontiers in Neuroinformatics*. 5.

19. Tustison NJ, Avants BB, Cook PA, Yuanjie Z, Egan A, Yushkevich PA, et al. (2010): N4ITK: Improved N3 Bias Correction. *IEEE Transactions on Medical Imaging*. 29:1310-1320.

20. Avants B, Epstein C, Grossman M, Gee J (2008): Symmetric diffeomorphic image registration with cross-correlation: Evaluating automated labeling of elderly and neurodegenerative brain. *Medical Image Analysis*. 12:26-41.

21. Zhang Y, Brady M, Smith S (2001): Segmentation of brain MR images through a hidden Markov random field model and the expectation-maximization algorithm. *IEEE Transactions on Medical Imaging*. 20:45-57.

22. Dale AM, Fischl B, Sereno MI (1999): Cortical Surface-Based Analysis. *NeuroImage*. 9:179-194.

23. Schneidman D, Klein A, Ghosh SS, Bao FS, Giard J, Häme Y, et al. (2017): Mindboggling morphometry of human brains. *PLOS Computational Biology*. 13.

24. Fonov VS, Evans AC, McKinstry RC, Almli CR, Collins DL (2009): Unbiased nonlinear average age-appropriate brain templates from birth to adulthood. *NeuroImage*. 47.

25. Evans AC, Janke AL, Collins DL, Baillet S (2012): Brain templates and atlases. *NeuroImage*. 62:911-922.

26. Wang S, Peterson DJ, Gatenby JC, Li W, Grabowski TJ, Madhyastha TM (2017): Evaluation of Field Map and Nonlinear Registration Methods for Correction of Susceptibility Artifacts in Diffusion MRI. *Frontiers in Neuroinformatics*. 11.

27. Huntenberg JM (2014): Evaluating nonlinear coregistration of BOLD EPI and T1w images [Master Thesis]: Freie Universität, Berlin.

28. Najbauer J, Treiber JM, White NS, Steed TC, Bartsch H, Holland D, et al. (2016): Characterization and Correction of Geometric Distortions in 814 Diffusion Weighted Images. *Plos One*. 11.

29. Jenkinson M, Bannister P, Brady M, Smith S (2002): Improved Optimization for the Robust and Accurate Linear Registration and Motion Correction of Brain Images. *NeuroImage*. 17:825-841.

30. Cox RW, Hyde JS (1997): Software tools for analysis and visualization of fMRI data. *NMR in Biomedicine*. 10:171-178.

31. Greve DN, Fischl B (2009): Accurate and robust brain image alignment using boundary-based registration. *NeuroImage*. 48:63-72.

32. Power JD, Mitra A, Laumann TO, Snyder AZ, Schlaggar BL, Petersen SE (2014): Methods to detect, characterize, and remove motion artifact in resting state fMRI. *NeuroImage*. 84:320-341.

33. Behzadi Y, Restom K, Liau J, Liu TT (2007): A component based noise correction method (CompCor) for BOLD and perfusion based fMRI. *NeuroImage*. 37:90-101.

34. Satterthwaite TD, Elliott MA, Gerraty RT, Ruparel K, Loughead J, Calkins ME, et al. (2013): An improved framework for confound regression and filtering for control of motion artifact in the preprocessing of resting-state functional connectivity data. *NeuroImage*. 64:240-256.

35. Patriat R, Reynolds RC, Birn RM (2017): An improved model of motion-related signal changes in fMRI. *NeuroImage*. 144:74-82.

36. Lanczos C (1964): Evaluation of Noisy Data. *Journal of the Society for Industrial and Applied Mathematics Series B Numerical Analysis*. 1:76-85.

37. Van Snellenberg JX, Slifstein M, Read C, Weber J, Thompson JL, Wager TD, et al. (2015): Dynamic shifts in brain network activation during supracapacity working memory task performance. *Human brain mapping*. 36:1245-1264.

38. Gordon EM, Laumann TO, Adeyemo B, Huckins JF, Kelley WM, Petersen SE (2016): Generation and Evaluation of a Cortical Area Parcellation from Resting-State Correlations. *Cerebral Cortex*. 26:288-303.

39. Van Essen DC, Curtiss SW, Laumann T, Jenkinson M, Prior F, Glasser MF, et al. (2011): Informatics and Data Mining Tools and Strategies for the Human Connectome Project. *Frontiers in Neuroinformatics*. 5.

40. Williams JC, Tubiolo PN, Luceno JR, Van Snellenberg JX (2022): Advancing motion denoising of multiband resting-state functional connectivity fMRI data. *NeuroImage*. 249.

41. Afyouni S, Nichols TE (2018): Insight and inference for DVARS. *NeuroImage*. 172:291-312.

42. Smyser CD, Snyder AZ, Neil JJ (2011): Functional connectivity MRI in infants: exploration of the functional organization of the developing brain. *NeuroImage*. 56:1437-1452.

43. Winkler AM, Webster MA, Vidaurre D, Nichols TE, Smith SM (2015): Multi-level block permutation. *NeuroImage*. 123:253-268.
